# The superficial cervical lymph nodes and mast cells contribute to sex-dependent cognitive impairment induced by a high-fat diet

**DOI:** 10.64898/2026.11.20.768228

**Authors:** Victor Manuel Ruiz-Rodríguez, Ares Orlando Cuellar-Santoyo, Angeles Braun-De Pedro, Karen Hernández-Balderas, Carlos Manuel Paredes-Popoca, Ana María Estrada-Sánchez

**Affiliations:** División de Biología Molecular, Laboratorio de Neurobiología, Instituto Potosino de Investigación Científica y Tecnológica (IPICYT), San Luis Potosí, San Luis Potosí, México

**Keywords:** Obesity, Neuroinflammation, Hippocampus, Chemokines, Cytokines, Mast cells, Periphery-brain axis

## Abstract

Obesity induces chronic inflammation linked to the development of multiple pathologies, including brain-related diseases. However, how peripheral inflammation from obesity increases brain vulnerability remains unclear. The superficial cervical lymph nodes and mast cells emerge as potential key contributors to brain vulnerability during peripheral inflammation. Therefore, we evaluated whether a high-fat diet (60% of calories from fat) induces peripheral inflammation and hippocampal neuroinflammation in male and female mice, impairs hippocampal function, and involves the superficial cervical lymph nodes. Our results showed that male mice exhibited short-term memory impairments after consuming a high-fat diet, whereas female mice were unaffected. Notably, only obese males showed increased expression of inflammatory markers (P2X1, P2X4, P2X7, NLRP3, IL-1β, ITGAL, CCR1, CCR5, CCR7) in superficial cervical lymph nodes, whereas the hippocampus showed only elevated *CCL3* mRNA levels, a chemokine produced by various immune cells, including mast cells, during neuroinflammation. When the mast cells in the hippocampus were evaluated, only obese male mice showed an increase in degranulated mast cells. To corroborate the contribution of the superficial cervical lymph nodes and mast cells to high-fat diet-induced short-term memory impairment, a lymphadenectomy was performed before the high-fat diet was initiated. Strikingly, mice subjected to lymphadenectomy gained no weight. Therefore, male mice showed no short-term memory impairment and exhibited fewer degranulated mast cells. Overall, male mice appear more susceptible to obesity-related cognitive deficits, likely driven by peripheral rather than neuroinflammatory mechanisms, with involvement of the superficial cervical lymph nodes and mast cells.

## 1 Introduction

Excessive body fat accumulation in obesity leads to low-grade, chronic systemic inflammation driven by cytokines and chemokines released by neutrophils, macrophages, and cytotoxic T lymphocytes that infiltrate adipose tissue (Alexaki, 2024; Le Thuc and Garcia-Caceres, 2024). This chronic peripheral inflammation significantly increases the risk of developing conditions such as type 2 diabetes, fatty liver disease, and cancer (Kawai et al., 2021). However, obesity is not only linked to peripheral organ disease but also to the development of brain disorders such as stroke and depression (Bluher, 2019). Thus, although the blood-brain barrier isolates the brain from peripheral alterations, it remains vulnerable to peripheral inflammation.

The glymphatic system serves as an interface between the periphery and the brain. It comprises the perivascular spaces surrounding the blood vessels that supply the brain and the lymphatic drainage system arising from the meninges and extending to the deep and superficial cervical lymph nodes (Louveau et al., 2015; Louveau et al., 2018; Rustenhoven et al., 2021; Le Thuc and Garcia-Caceres, 2024; Jin et al., 2025; Zhao et al., 2026). The deep cervical lymph nodes are a major drainage route for meningeal lymphatics (Louveau et al., 2018; Rustenhoven et al., 2021), but the cervical lymphatic system comprises interconnected, functionally related drainage pathways rather than entirely independent compartments (Louveau et al., 2015; Louveau et al., 2018). In this sense, emerging evidence shows that superficial cervical lymph nodes also contribute to brain drainage, but less is known about their contribution to the pathogenesis of brain diseases (Jin et al., 2025). Both deep and superficial cervical lymph nodes contain various immune cells that facilitate signaling between the brain and the periphery, enable recognition of antigens originating in the brain, recruit and activate T lymphocytes in the meninges, and promote the activation of mast cells that can migrate from the periphery to the brain (Silverman et al., 2000; Louveau et al., 2018; Rustenhoven et al., 2021; Elieh-Ali-Komi et al., 2025).

In addition to the cervical lymph nodes, mast cells are present in peripheral tissues and the brain parenchyma, particularly on the abluminal side of blood vessels, where they interact with neurons, glial cells, and endothelial cells (Hendrikus et al., 2017). During obesity, mast cells release cytokines (e.g., IL-1β, TNF-α) and chemokines (e.g., CCL2, CCL3) that promote inflammation in adipose tissue and, in the brain, activate microglia (Shi and Shi, 2012; Tian et al., 2025). Activated mast cells release granules containing a variety of mediators, including cytokines, chemokines, proteases, neurotransmitters, and reactive oxygen species (Theoharides et al., 2019; Aitella et al., 2026; Derakhshani et al., 2026). Their overactivation is also linked to neurodegenerative processes and neuropathologies (Hendrikus et al., 2017). Therefore, mast cells might be a key contributor to peripheral inflammation and brain vulnerability during obesity.

Obesity-related alterations are sex-dependent, driven by differences in immune responses, visceral fat accumulation, inflammation, and metabolic changes, all of which are regulated by sex hormones, sex chromosome-specific gene expression, and epigenetic mechanisms (Shepherd et al., 2020; Vijayakumar et al., 2026). Thus, its impact on the brain also differs between male and female mice, particularly in the hippocampus, a brain region vulnerable to obesity (Baghdadchi et al., 2026). However, it remains unknown whether communication mediated by superficial cervical lymph nodes between the periphery and the brain contributes to sex-related differences in obesity-associated hippocampal function. Therefore, this study aimed to determine whether superficial cervical lymph nodes are key to changes in hippocampal function during obesity and whether these changes are related to mast cells in male and female mice.

## 2 Materials and Methods

### 2.1 **Mice and diets**

Three-month-old male and female C57BL/6 mice were used in this study. The experiments presented were obtained from different cohorts of mice, including males and females from the same litter, resulting in an imbalance in the number of male and female mice used. All mice were individually housed in polypropylene cages under a 12-h light/dark cycle with ad libitum access to food and water. Based on previous studies (Sanchez-Tapia et al., 2024; Totten et al., 2025), mice were divided into a control group fed a standard diet (LabDiet, 5001; 13.5% calories from fat, 58% from carbohydrates, and 28.5% from protein) and purified water, and an experimental group fed a high-fat diet (60% calories from fat, 27% from carbohydrates, and 13% from protein) with 5% sucrose in the drinking water for 16 weeks. During this period, body weight, blood glucose, cholesterol, and triglyceride levels were measured. A mouse was considered obese if it exhibited a significant increase in visceral fat accumulation compared to the control group, as previously described (Mandarim-de-Lacerda et al., 2021). All experimental procedures were conducted in accordance with the National legislation for the care and use of experimental animals (Consejo Mexicano de Cuidado Animal, Norma Oficial Mexicana NOM-062-ZOO-1999). All procedures were approved by the committee for experimental use of animals of the Instituto Potosino de Investigación Científica y Tecnológica (IPICYT) under protocol number 18c.

### 2.2 Glucose and insulin tolerance tests

Glucose tolerance and insulin tolerance tests were performed on 11 mice (4 females and 7 males) fed a control diet and on 11 mice (4 females and 7 males) fed a high-fat diet. For this test, blood was obtained from the tail after placing the mice in a restraint chamber that allowed the tail to be exposed. We collected blood by massaging the tail and then making a single pinch at the base of the tail. Glucose levels were measured using the Accu-Chek Active Glucose Meter (Roche). The glucose tolerance test was performed after 12 hours of fasting. Initial basal glucose was measured, followed by measurements at 15, 30, 60, and 120 minutes after an intraperitoneal injection of 2 g/kg D-(+)-glucose (Sigma-Aldrich) in 0.9% saline. For the insulin tolerance test, the mice were fasted for 4 hours. Blood glucose was measured at baseline and 15, 30, 60, and 120 minutes after an intraperitoneal injection of 0.9 U/kg of rapid-acting insulin diluted in 0.9% saline solution.

### 2.3 Novel object recognition test

We used the novel object recognition test to evaluate short-term memory, as previously described by (Moore et al., 2013; Hauser et al., 2017; Cuellar-Santoyo et al., 2026). Briefly, mice were placed in a transparent acrylic open box (50 x 50 x 30 cm) in a noise-free room with controlled environmental conditions (22°C under red light). The test was divided into three consecutive phases. Initially, the mouse explored the box freely for 5 minutes, then was removed and placed in its home cage for about one minute. In the second phase, three objects of the same color, shape, and texture were placed on the left side of the box. The mice were then returned and allowed to explore the box and the identical objects for 10 minutes. After this period, the mice were removed and returned to their home cage for two minutes (Moore et al., 2013). In the final phase, the central object was replaced with another of a different color, shape, and rough texture, and the mice were allowed to explore freely for an additional 10 minutes. The last two phases of the behavioral tests were recorded on video for later analysis. To minimize bias, the video analysis was conducted by an experimenter blinded to the identities of the experimental groups. For the novel object recognition test, 29 mice were used: 13 (6 females and 7 males) fed a control diet, and 16 (8 females and 8 males) fed a high-fat diet. After finishing the behavioral task, brain samples were collected for the various molecular and imaging assays described below.

Based on the results, mice’s performance on the novel object recognition test was assessed by calculating the exploration preference (EP) and the discrimination index, two complementary measures of novel-object exploration, using the following formulas for EP for the familiar object (FO) or the novel object (NO), as previously described by Cuellar-Satoyo 2026 (22):

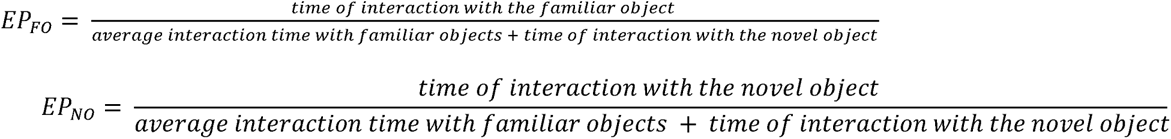

The discrimination index was computed using the formula:

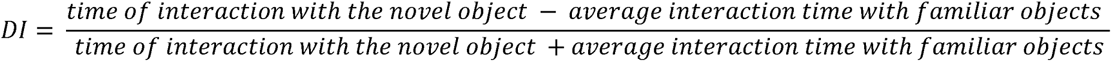

### 2.4 RT-qPCR

Mice were anesthetized with sodium pentobarbital (100 mg/kg) and perfused with cold 0.9% saline to remove blood. The hippocampus and lymph nodes were then dissected and immediately frozen in liquid nitrogen. Each frozen sample was mechanically pulverized, and total RNA was extracted using TRIzol and a phenol-chloroform gradient. Complementary deoxyribonucleic acid (cDNA) was synthesized from 1 µg of total RNA in 10 µL using the high-capacity reverse transcription kit (Thermo Scientific, Massachusetts, USA) according to the manufacturer’s recommendations. The cDNA concentration for all samples was adjusted to 200 ng/μL. Real-time PCR was performed on a CFX96 real-time system (Bio-Rad, CA, USA). We used 10 μL reaction volumes containing 2.5 μL cDNA, 5 μL Maxima SYBR Green Master Mix 2x (Thermo Scientific, Massachusetts, USA), 0.3 μL of each primer (300 nM), and 1.9 μL of H_2_O DEPC. The primer sequences used to identify the different inflammatory markers are shown in **Table 1**. Results were obtained by normalizing each sample to its endogenous control (β-actin or 18S) and using the 2^-ΔΔCt method to calculate fold changes relative to the control group or the female mouse group. For the hippocampus, both hemispheres were processed separately, and each sample was run in duplicate. The fold change for each hemisphere was calculated individually, and the values were averaged to obtain a single value per animal. For this experiment, 11 control mice (5 females and 6 males) and 11 mice on a high-fat diet (5 females and 6 males) were used.

**Table 1.**
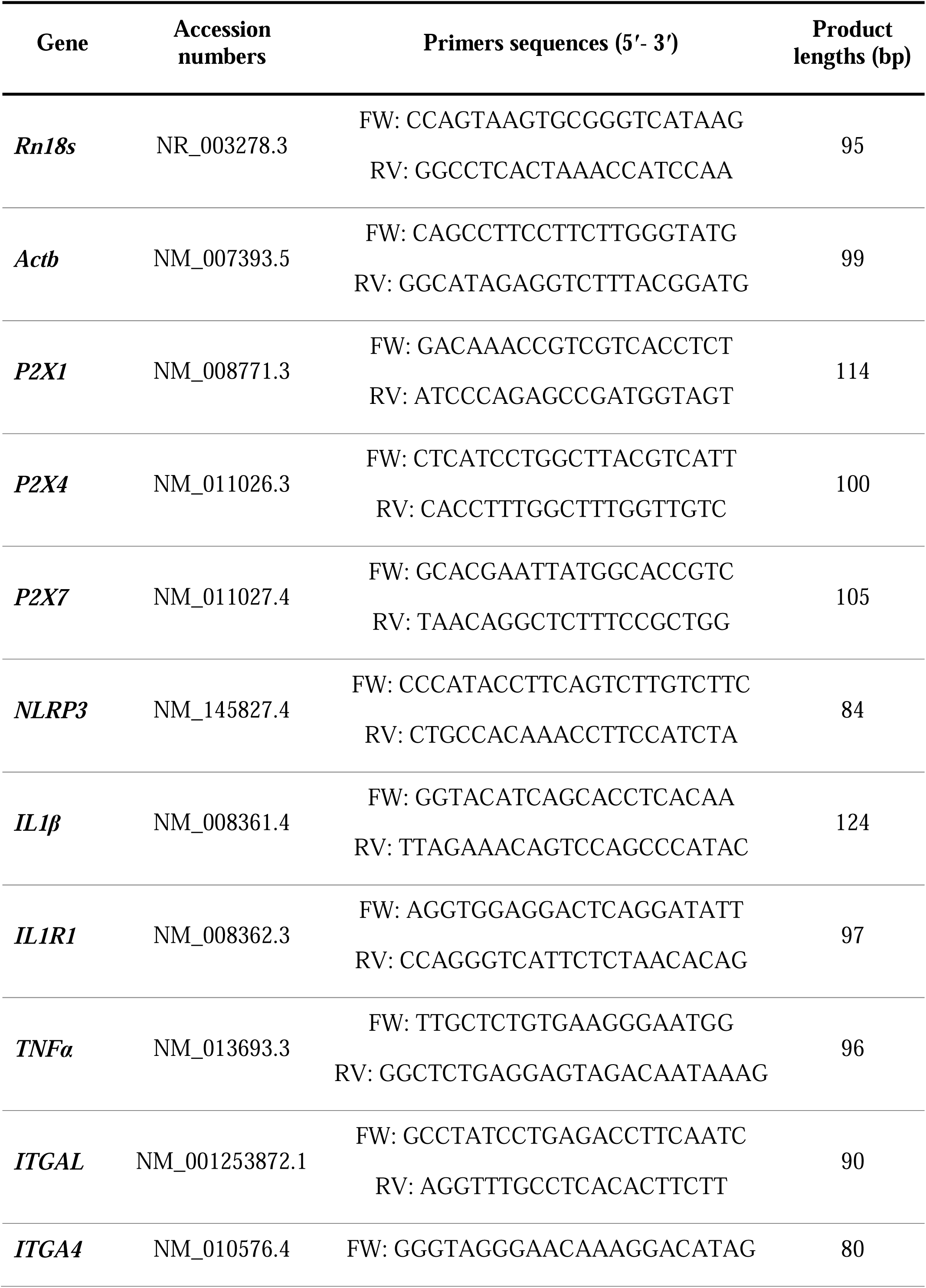
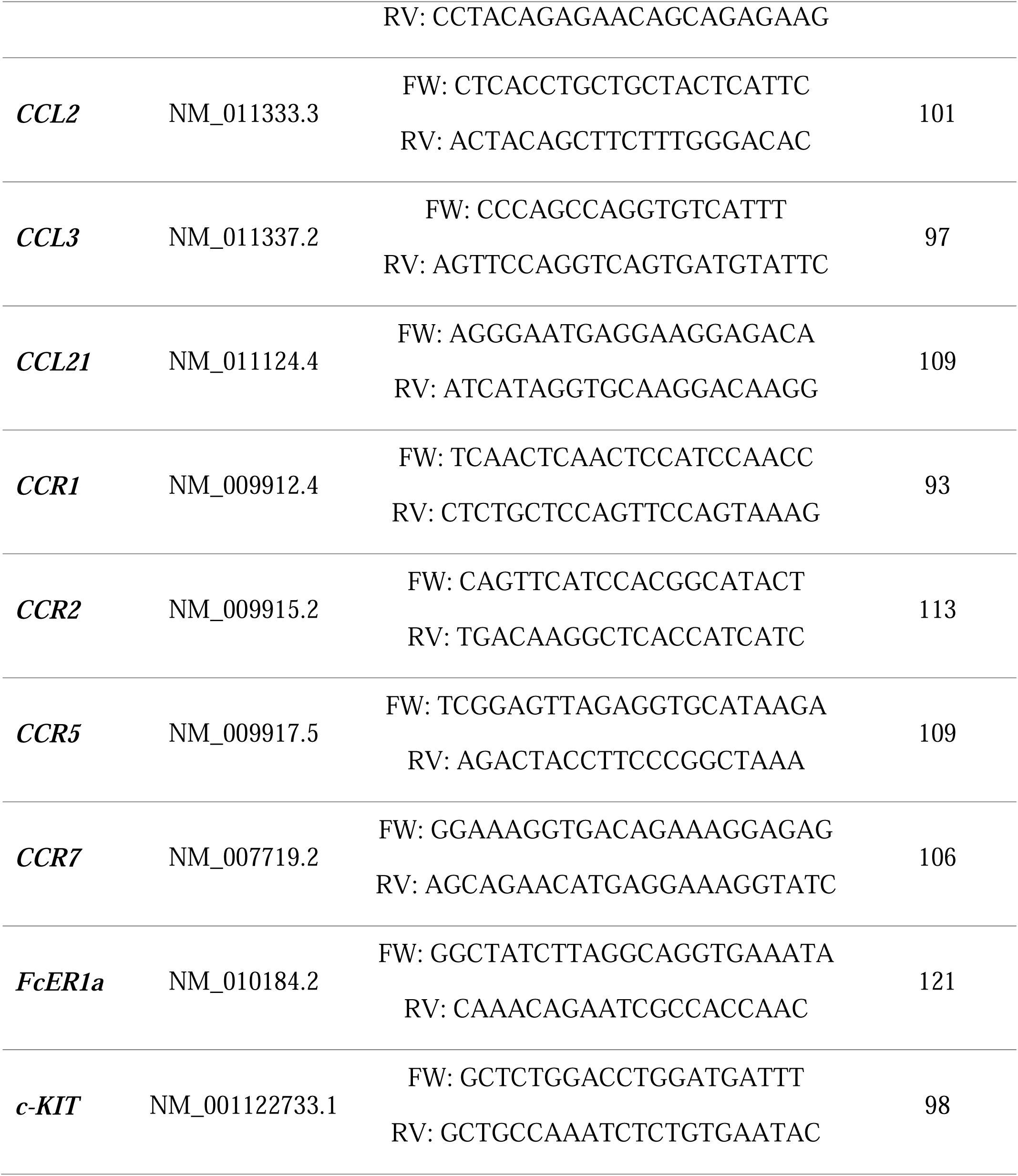
Primers sequences used for RT-qPCR analysis.

### 2.5 Mast cell staining and quantification

Mast cells were visualized by toluidine blue staining, as described by Nautiyal 2012 (Nautiyal et al., 2012). Briefly, 5-µm sections were obtained from paraffin-embedded superficial cervical and inguinal lymph nodes and brains using an automatic microtome (SLEE, 6062). The slice-mounted sections were deparaffinized in xylene and rehydrated using an ethanol-to-water gradient. The slices were immersed in toluidine blue solution (0.5% toluidine blue in 60% ethanol, pH 2.3) for 10 min. The sections were then briefly rinsed in distilled water, dehydrated with a series of ethanol washes (2 × 95% EtOH for 1 min, 2 × 100% EtOH for 1 min) and xylene (2 × 100% for 30 s), and mounted with Entellan (Merck). The total number of mast cells was quantified, and they were subsequently classified. A mast cell was considered granular when its cytoplasm was filled with tightly packed metachromatic granules that could not be distinguished individually, the purple staining was intense and uniform, and the outline was sharp. A mast cell was considered degranulated when metachromatic granules were visible outside the cell outline, or when it stained a paler purple than neighboring mast cells in the same section, usually with individual granules distinguishable in the cytoplasm, as previously described by different authors (Larson et al., 2011; Lenz et al., 2018; Dong et al., 2019). For this assessment, 6 control mice (3 females and 3 males) and 8 mice (4 females and 4 males) fed a high-fat diet were used. In addition, for the superficial cervical lymphadenectomy group, we used 6 mice (3 females and 3 males) on a control diet and 6 mice (3 females and 3 males) on a high-fat diet. For each mouse, a coronal section of the left and right lymph nodes and two coronal sections of the hippocampus were analyzed. Whole sections of superficial cervical lymph nodes were reconstructed from 5× images (1.29 µm/pixel). Adjacent 5× fields were aligned using SIFT features (Lowe, 2004) and merged when at least 25 geometrically consistent matches were found. Each 20× field was downsampled to 5× resolution and placed in its 5× mosaic via normalized cross-correlation (Muhlich et al., 2022). Mast cells were detected in 20× fields using a Cellpose-SAM model (Pachitariu et al., 2025) fine-tuned on our images. We tested the model on 39 images that we manually counted, without knowing its output. Detected cells were mapped to mosaic coordinates. Where 20× fields overlapped, cells less than 10 µm apart were counted once. Clustering was evaluated with the Clark–Evans nearest-neighbor index (Clark and Evans, 1954). The observed value was compared with 999 simulations in which cells were placed at random within the same tissue. Each simulation preserved the observed number of cells inside and outside the parenchyma. A hotspot was defined as any 250 µm grid square whose mast cell count exceeded the 95th percentile of the per-simulation maximum square count. Using the maximum controls for multiple comparisons across squares. We produced local density maps with a Gaussian kernel (σ = 100 µm), as is common in spatial analyses of immune cells in tissue (Page et al., 2023). The image-analysis code (Python 3; OpenCV, scikit-image, SciPy and statsmodels) was written with the help of Claude (Anthropic). The authors set the analysis criteria and checked each step: model detections were validated against blind manual counts, and the results were reviewed on annotated images. Manual counts and review of regions of interest were done in Fiji.

### 2.6 CCL3 immunohistochemistry

For mast cell staining and CCL3 co-detection, we used the immunohistochemistry protocol described by (Mares-Barbosa et al., 2025). Briefly, the hippocampus and the superficial cervical lymph nodes were embedded in paraffin, and 5 µm coronal sections were obtained using a microtome (SLEE, 6062 automatic). We used the Pierce™ diamino-benzidine substrate kit (Thermo Fisher Scientific) to analyze CCL3 chemokine content by immunohistochemistry. Sections were deparaffinized in xylene and rehydrated using an ethanol-to-water gradient. Sections were then treated with Tris-EDTA buffer (pH 9.0) in a pressure cooker for antigen retrieval. Samples were treated with 3 % H_2_O_2_ to inactivate endogenous peroxidases. Sections were incubated overnight at 4°C with a goat polyclonal CCL3 antibody (#79554347 from Thermo Fisher Scientific) at a 1:25 dilution in a humid chamber. Sections were washed and incubated with a Peroxidase AffiniPure Donkey Anti-Goat IgG secondary antibody (Jackson Immuno Research) at a 1:500 dilution for two hours at room temperature. Samples were treated with diaminobenzidine substrate for 10 minutes, washed twice with phosphate-buffered saline and Tween solution, and mounted using Aquatex® mounting medium (Merck). Images were taken with the Axio Imager.M2 microscope (ZEISS).

### 2.7 Cervical lymphadenectomy

In a separate cohort, 6 mice (3 females and 3 males) on a control diet and 9 (5 females and 4 males) on a high-fat diet underwent bilateral superficial cervical lymphadenectomy. Briefly, the mice were anesthetized with ketamine (90 mg/kg), xylazine (10 mg/kg), and acepromazine (2 mg/kg) in sterile saline and administered at 0.1 mL per 10 g of body weight. The surgery began only after the pedal withdrawal reflex was lost. Their necks were shaved, and the skin was disinfected with iodine and 70% ethanol. A 5-mm midline skin incision was made to expose the submandibular salivary glands; three superficial cervical lymph nodes of the mandibular group were identified on each side (Fig. 8A). Two were located at the rostral border of the submandibular gland, just caudal to the masseter muscle, and one was located along the lateral border of the submandibular gland, near the junction of the facial veins, which became visible after the gland was gently retracted. We freed the lymph nodes from surrounding adipose and connective tissue with fine forceps and removed them. The skin was closed with Histoacryl tissue adhesive (Braun). In sham surgery (NS mice), the same skin incision was made and then sutured without exposing or removing the superficial cervical lymph nodes. Five days after surgery, they started on the corresponding diet.

### 2.8 Statistical analysis

All data were analyzed using GraphPad Prism version 10.2.2 (GraphPad Software Inc., San Diego, CA, USA). Data were first evaluated to determine whether they followed a parametric or nonparametric distribution, and the appropriate statistical analysis was then performed. Differences between mice fed the control diet and those fed the high-fat diet were assessed using an unpaired t-test or the Mann-Whitney U test. To evaluate the diet’s effects on the genes evaluated, we first compared the groups, then evaluated effects by sex and diet using two-way ANOVA followed by a post hoc test. To examine sex-dependent differences, multiple Mann-Whitney tests and two- or three-way ANOVAs were used, followed by post hoc tests (Bonferroni, Tukey, or Fisher’s). Glucose tolerance and insulin tolerance data were analyzed using a two-way repeated-measures ANOVA with Bonferroni post hoc testing. Correlation matrices were calculated using Spearman’s correlation coefficient. All data are expressed as mean ± SEM; differences were considered statistically significant at *p* < 0.05.

## 3 Results

### 3.1 **The consumption of a high-fat diet for 16 weeks causes obesity in female and male mice**

Average food intake, weight gain, and visceral fat accumulation were evaluated at the end of the 16-week high-fat diet. Mice fed the high-fat diet consumed less food than those in the control group (3.17 ± 0.30 vs. 4.78 ± 0.53 g/day; *p* < 0.0001). This reduction was observed in both sexes: females on the high-fat diet consumed 3.28 ± 0.25 g/day, compared with 4.49 ± 0.50 g/day in the control group (*p* = 0.020), and males in the control group consumed 3.09 ± 0.34 g/day, compared with 4.99 ± 0.50 g/day in those fed the high-fat diet (*p* = 0.0008). Overall, mice fed the high-fat diet gained significant weight (Fig. 1A; *p* < 0.0001), and this effect was observed by sex: female mice (*p* = 0.0202) and male mice (*p* = 0.0090; Fig. 1B), but with no interaction between the variables of diet and sex (*p* = 0.9724). Likewise, the high-fat diet-fed group showed increased visceral fat mass (Fig. 1C; *p* < 0.0001), an effect that persisted when comparing female (*p* = 0.0149) and male (*p* = 0.0138) mice to their respective control groups (Fig. 1D; interaction diet/sex *p* = 0.09201). Male and female mice gained weight and accumulated visceral fat similarly on the high-fat diet (Figs. 1B and 1D).

**Figure 1.**
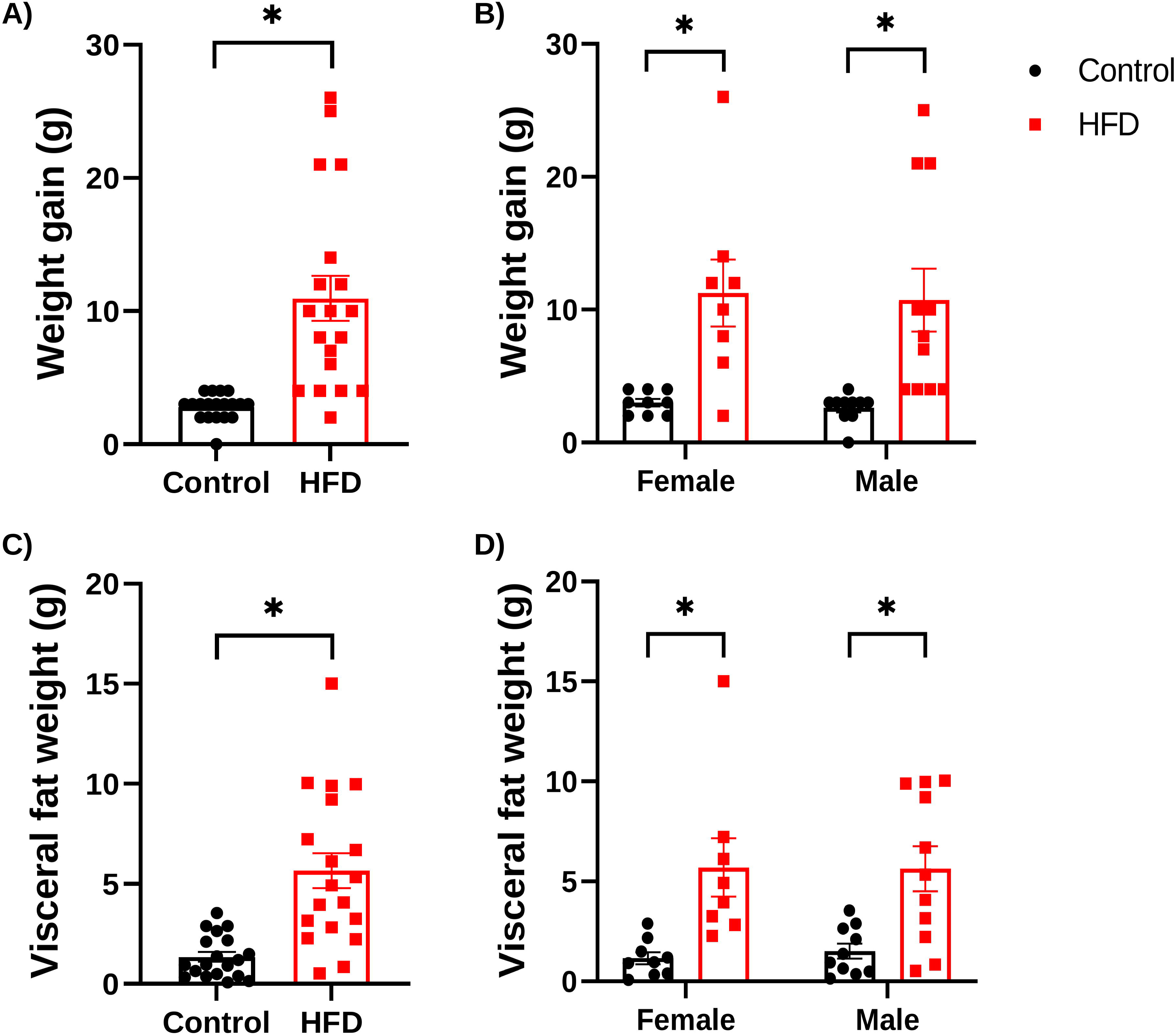
Weight gain and visceral fat accumulation in female and male mice fed a control diet or a high-fat diet. The graphs show weight gain (A and B) and visceral fat weight (C and D) in female and male mice fed either a control or a high-fat diet for 16 weeks. Data were analyzed using the Mann-Whitney test to compare the control and high-fat diet groups. To assess differences between sexes, multiple Mann-Whitney tests or a two-way ANOVA with Bonferroni post hoc testing were performed. Data are presented as mean ± SEM (* p < 0.05).

### 3.2 Altered metabolic parameters in female and male mice fed a high-fat diet

Next, we evaluated whether the high-fat diet affected glucose, insulin, or cholesterol levels. Eight weeks after starting the high-fat diet, the glucose tolerance test showed that mice had higher blood glucose levels 30 minutes after glucose injection (*p* = 0.0104). At 16 weeks, mice fed a high-fat diet had higher glucose levels at 15 (*p* = 0.0327) and 30 minutes (*p* = 0.0295) after glucose injection (Fig. 2A). Thus, mice fed a high-fat diet exhibited impaired glucose tolerance compared with controls.

**Figure 2.**
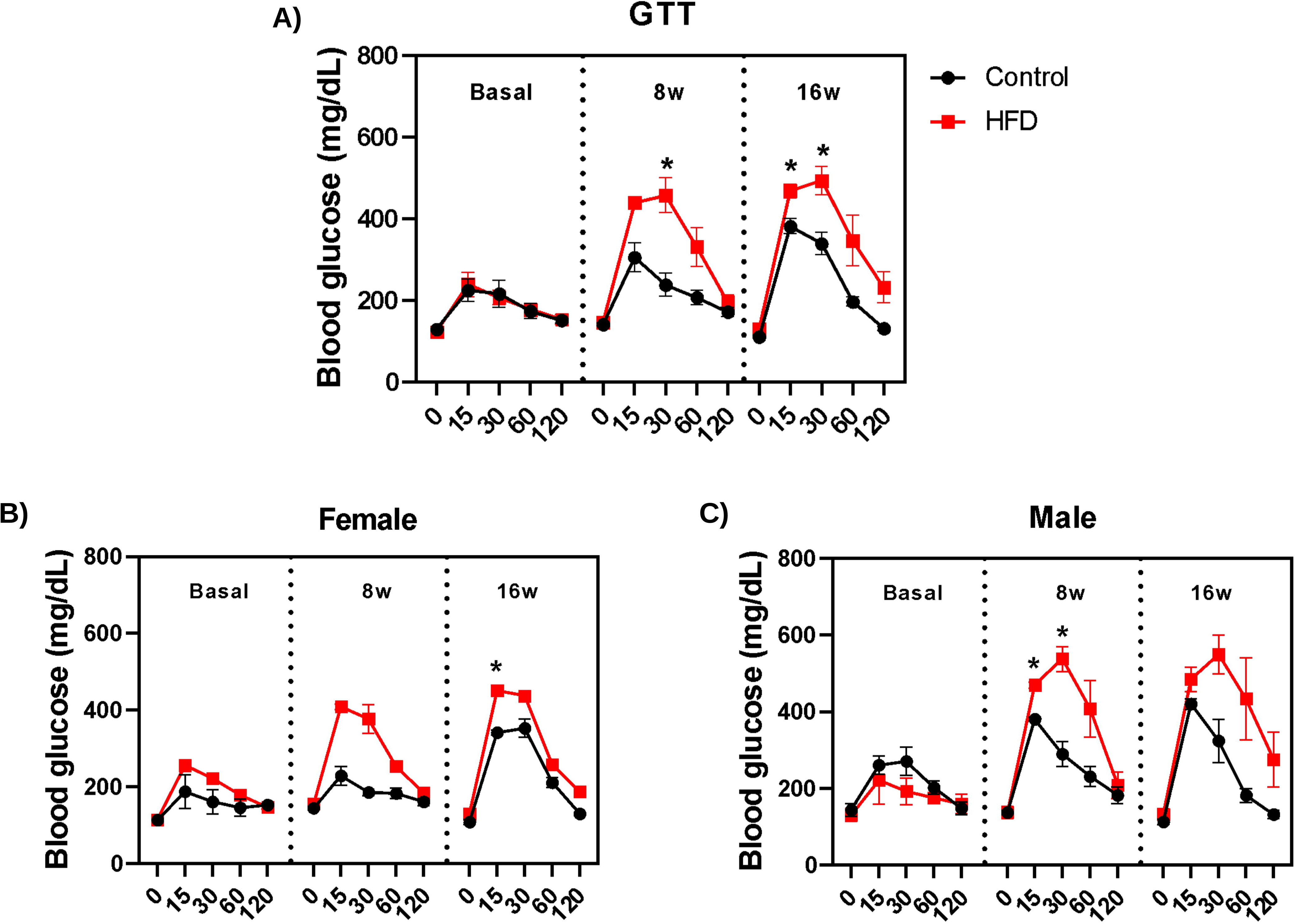
Glucose tolerance test (GTT) in female and male mice fed a control diet or a high-fat diet. (A) Blood glucose kinetics in mice at the start of the experiment (basal) and at 8 and 16 weeks (8w and 16w) after being fed a control diet or a high-fat diet. Differences in blood glucose kinetics in female (B) and male (C) mice fed a control diet or a high-fat diet. Data were analyzed using a two-way repeated-measures ANOVA with Bonferroni post hoc testing. Data are presented as mean ± SEM (* p < 0.05).

When evaluating the effect of sex on glucose levels, female mice showed a significant increase in blood glucose concentration only at 16 weeks on a high-fat diet, 15 minutes after glucose injection (Fig. 2B; *p* = 0.0062). In male mice, altered glucose tolerance was observed earlier, at 8 weeks of high-fat diet consumption (Fig. 2C), at 15 minutes (*p* = 0.0055) and at 30 minutes (*p* = 0.0308) after glucose injection.

In contrast to the glucose tolerance test results, no insulin resistance was observed in mice fed a high-fat diet (Supplementary Fig. 1A-C; *p* > 0.9999 in all analyses).

Likewise, high-fat diet consumption did not affect blood triglyceride levels (Fig. 3A and B; *p* > 0.1 in all analyses; interaction diet/sex *p* = 0.6177). However, blood cholesterol levels increased significantly in mice fed a high-fat diet compared with the control group (Fig. 3C; *p* < 0.0001). This effect was also observed by sex; female (*p* = 0.0004) and male (*p* = 0.0004) mice fed a high-fat diet showed increased blood cholesterol (Fig. 3D; interaction diet/sex *p* = 0.5384).

**Figure 3.**
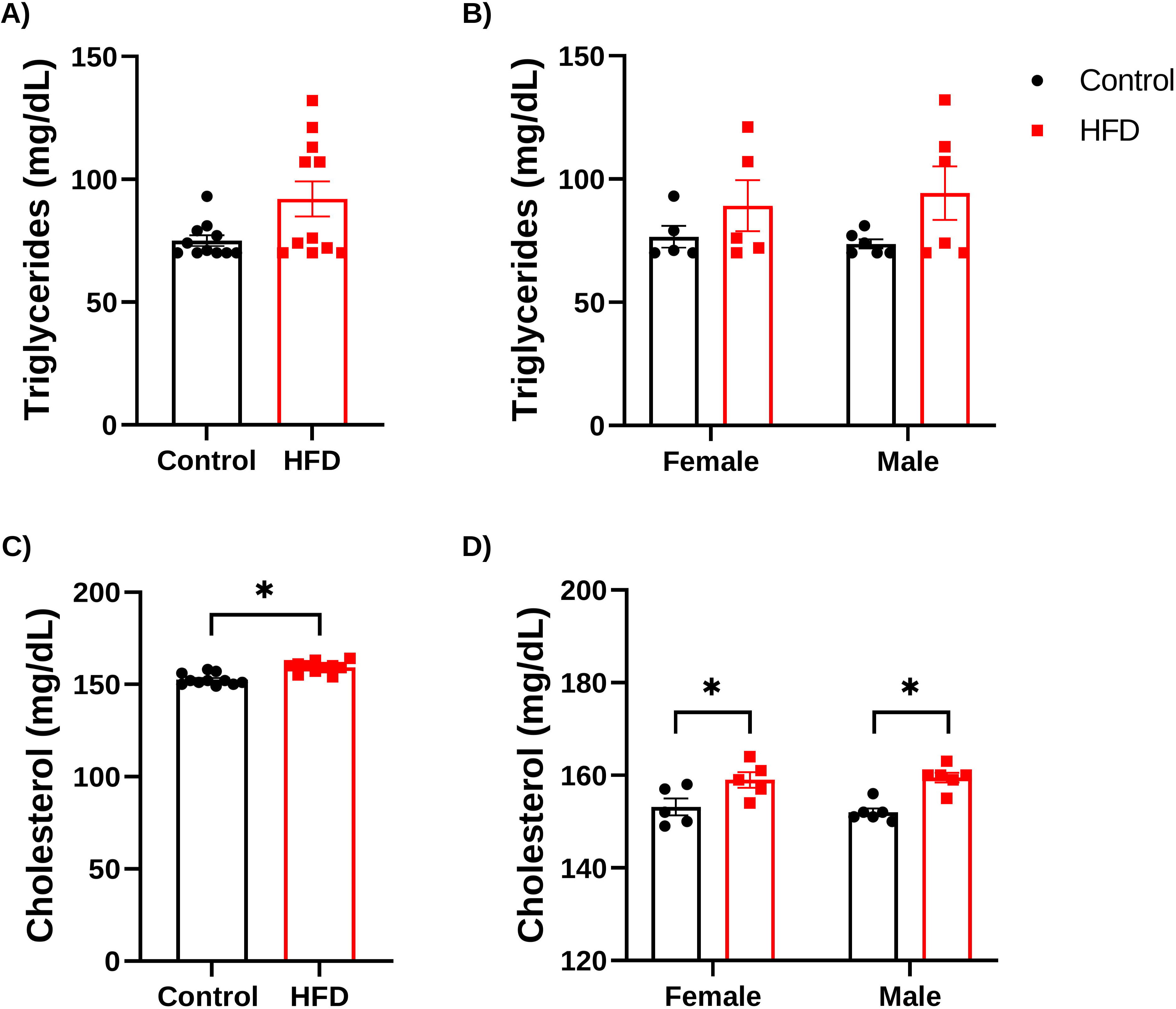
Blood triglyceride and cholesterol levels in mice fed a control diet or a high-fat diet. The graphs show blood triglyceride (A and B) and cholesterol (C and D) levels in female and male mice fed either a control diet or a high-fat diet for 16 weeks. Data were analyzed using the Mann-Whitney test or an unpaired t-test to assess differences between the control and high-fat diet groups. To assess differences between sexes, a two-way ANOVA was performed, followed by a Bonferroni post hoc test. Data are presented as mean ± SEM (* p < 0.05).

### 3.3 Consumption of a high-fat diet impairs short-term memory only in male mice

We assessed short-term memory using the novel object recognition test to evaluate hippocampal function after the high-fat diet. Total exploration time for control and mice fed the high-fat diet is shown in Supplementary Fig. 2. Control mice spent more time exploring the novel object than the familiar object (Supplementary Fig. 2A; *p* < 0.0001), whereas mice fed the high-fat diet explored both objects for a similar amount of time (*p* > 0.9999) and spent less time with the novel object than control mice (*p* < 0.0001). Accordingly, control mice preferred to explore the novel object (Fig. 4A; *p* < 0.0001), whereas mice fed the high-fat diet showed a lower preference for the novel object (*p* = 0.0164). By sex, female control mice spent more time exploring the novel object than the familiar object (Supplementary Fig. 2B; *p* = 0.0332), whereas the high-fat diet group did not (*p* > 0.9999). Female control and high-fat diet groups showed similar exploration preferences for the novel object (Fig. 4B; *p* > 0.9999). In contrast, male control mice spent more time with the novel object (Supplementary Fig. 2C; *p* < 0.0001), therefore preferentially explored the novel object (Fig. 4C; *p* < 0.0001), but the high-fat diet group showed a significant reduction in exploration time (*p* < 0.0001) and in preference for the novel object (*p* = 0.0076).

**Figure 4.**
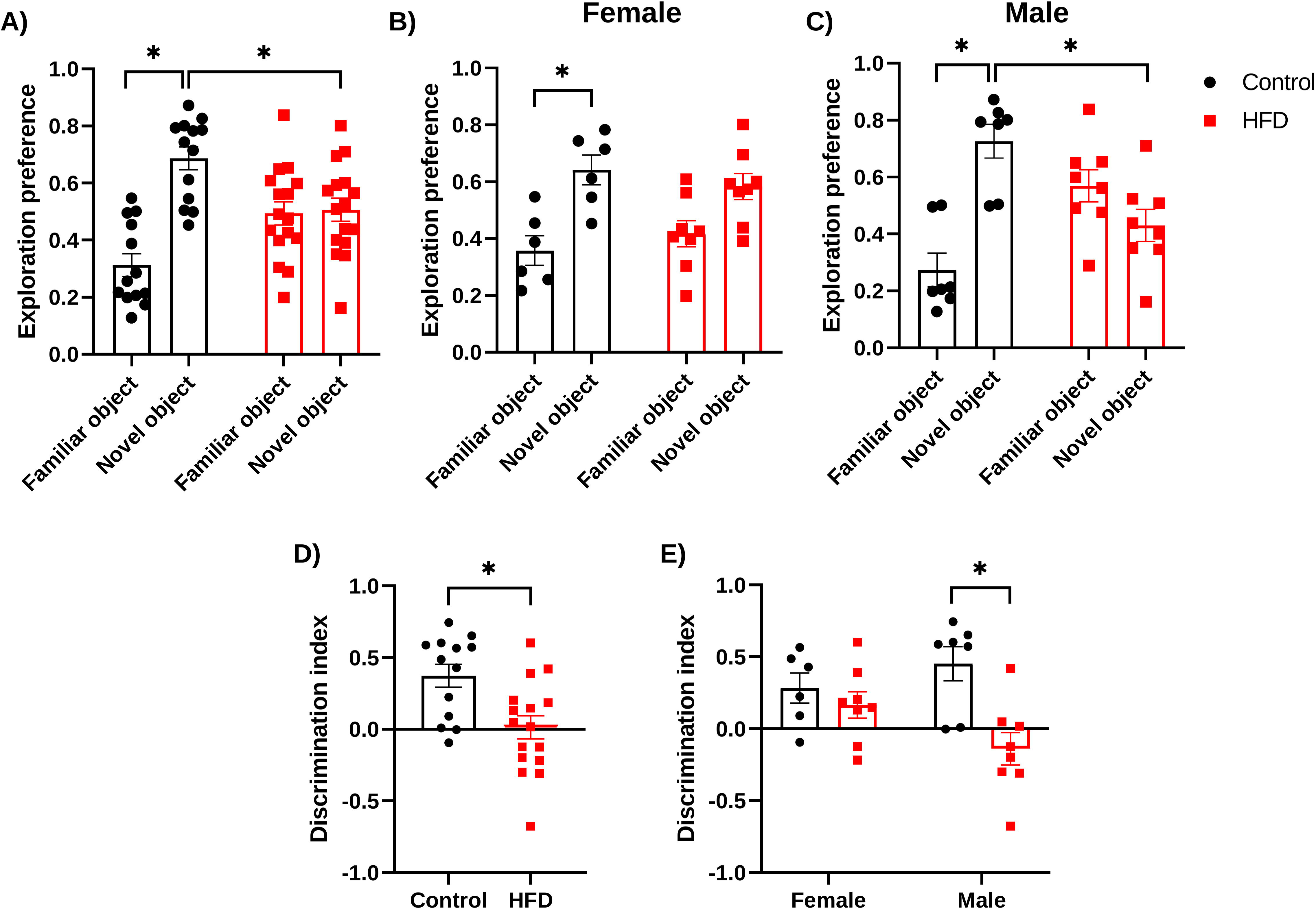
Novel object recognition test in female and male mice fed a control diet or a high-fat diet. (A) Exploration preference for the familiar and novel objects in mice fed a control diet or a high-fat diet for 16 weeks. The graphs show changes in exploration preference in female (B) and male (C) mice. (D) Discrimination index in mice fed a control diet or a high-fat diet for 16 weeks. (E) Differences in the discrimination index between female and male mice. Data were analyzed using an unpaired t-test and two-way ANOVA with Bonferroni post hoc testing. Data are presented as mean ± SEM (* p < 0.05).

Mice fed a high-fat diet had a lower discrimination index than the control group (Fig. 4D; *p* = 0.0040). We also observed a significant diet × sex interaction (p = 0.0387). When comparing by sex, only male mice had a lower discrimination index than their control group (Fig. 4E; *p* = 0.0034).

Next, we evaluated velocity as an index of movement because weight gain might affect mice’s exploration of the objects. Overall locomotion speed analysis revealed a mean velocity of 15.87 ± 0.79 cm/s in the control group and 13.51 ± 1.03 cm/s in the high-fat diet group (p = 0.0686, Mann– Whitney U test). When stratified by sex, control males exhibited a mean velocity of 15.08 ± 1.59 cm/s, compared with 13.97 ± 1.75 cm/s in high-fat diet-fed males (p > 0.9999). Similarly, control females displayed a mean velocity of 16.67 ± 0.00 cm/s, whereas high-fat diet-fed females reached 13.06 ± 1.23 cm/s (p = 0.0606, Mann–Whitney U test). Therefore, body weight does not appear to affect exploration of the familiar or novel objects during the test.

Overall, these results indicate that male mice showed impaired short-term memory after consuming the high-fat diet, suggesting that the hippocampus of these mice is more vulnerable to the harmful effects of a high-fat diet, an effect that was not observed in female mice.

### 3.4 A high-fat diet increased the relative mRNA expression of only the chemokine *CCL3* in the hippocampus

To determine whether neuroinflammation underlies high-fat diet-induced impairment of hippocampal function in males, we measured the relative expression of various inflammatory mediators (Fig. 5). These included the purinergic receptors *P2X1*, *P2X4*, and *P2X7*; the *NLRP3* inflammasome component; *IL-1*β and its receptor *IL-1R1*; *TNF-*α; the chemokines *CCL2*, *CCL3*, and *CCL21*; and mast cell markers such as *FcER1*α and *c-KIT*. Overall, most inflammatory markers in mice fed a high-fat diet did not differ from those in the control group (*p* > 0.1); the same was observed when the data were analyzed by sex (Fig. 5A-Q and S-X; *p* > 0.1). However, only the chemokine *CCL3* showed a significant increase in expression in male mice fed a high-fat diet compared with their control group (Fig. 5R; *p* = 0.0330). Female mice showed similar levels between the control and high-fat diet groups (*p* = 0.0727). The data also showed a significant interaction between diet and sex (*p* = 0.0083). Because we observed increased relative *CCL3* expression in the hippocampus, we performed immunodetection of this chemokine in the hippocampus and found no statistically significant differences (data not shown).

**Figure 5.**
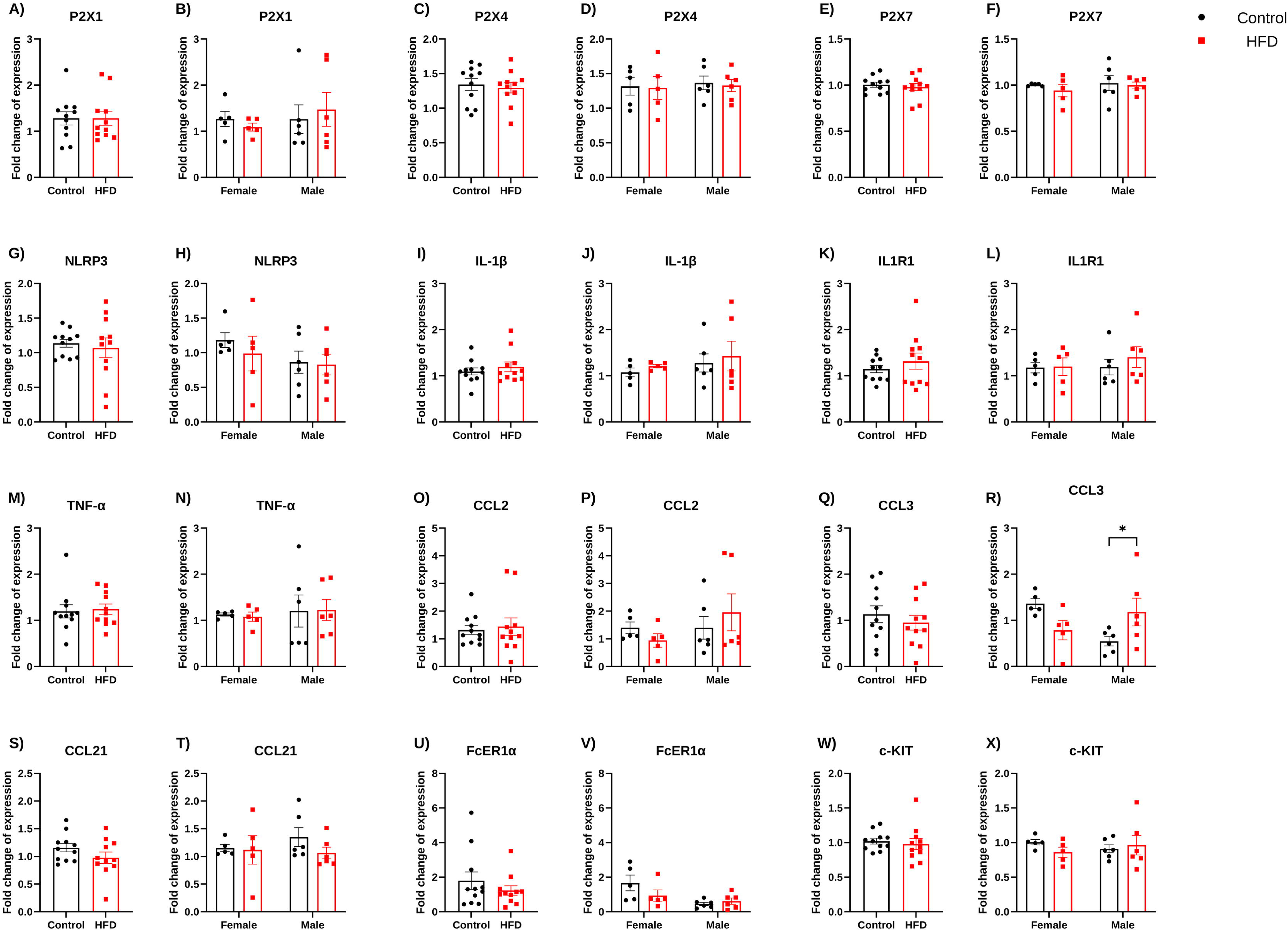
Relative mRNA expression levels of inflammatory mediators in the hippocampus of mice fed a control diet or a high-fat diet. The graphs show relative mRNA expression in the hippocampus of mice fed a control or high-fat diet for 16 weeks, with sex differences for genes encoding the purinergic receptors *P2X1* (A and B), *P2X4* (C and D), and *P2X7* (E and F); the *NLRP3* inflammasome component (G and H); *IL-1*β (I and J) and its receptor *IL-1R1* (K and L); *TNF-*α (M and N); the chemokines *CCL2* (O and P), *CCL3* (Q and R), and *CCL21* (S and T); and the mast cell markers *FcER1a* (U and V) and *c-KIT* (W and X). Data were analyzed using the unpaired t-test or the Mann-Whitney test to assess differences between the control and high-fat diet groups. To evaluate sex-dependent differences between groups, multiple Mann-Whitney tests or two-way ANOVA with Fisher’s LSD post hoc test were used. Data are presented as mean ± SEM (* p < 0.05). Each point represents one animal (mean of the left and right hippocampus).

### 3.5 Male mice fed a high-fat diet had an increased number of mast cells and mast cell degranulation in regions adjacent to the hippocampus

CCL3 can be released by various cells in the brain, but mast cells, a type of immune cell located in the brain parenchyma and the periphery, have been described as contributing to neuropathological processes (Shi and Shi, 2012; Tian et al., 2025; Ruiz-Rodriguez et al., 2026). Thus, we quantified granulated and degranulated mast cells in the hippocampus and adjacent regions (lateral ventricle, third ventricle, cortex, and corpus callosum overlying the hippocampus; Fig. 6F). Overall, the high-fat diet group had more mast cells (Fig. 6B; *p* = 0.0444). By sex, male mice fed a high-fat diet had more mast cells than their control group (Fig. 6C; *p* = 0.0059) and than female mice fed a high-fat diet (*p* = 0.0138). The data also showed a significant interaction between diet and sex (*p* = 0.0040).

**Figure 6.**
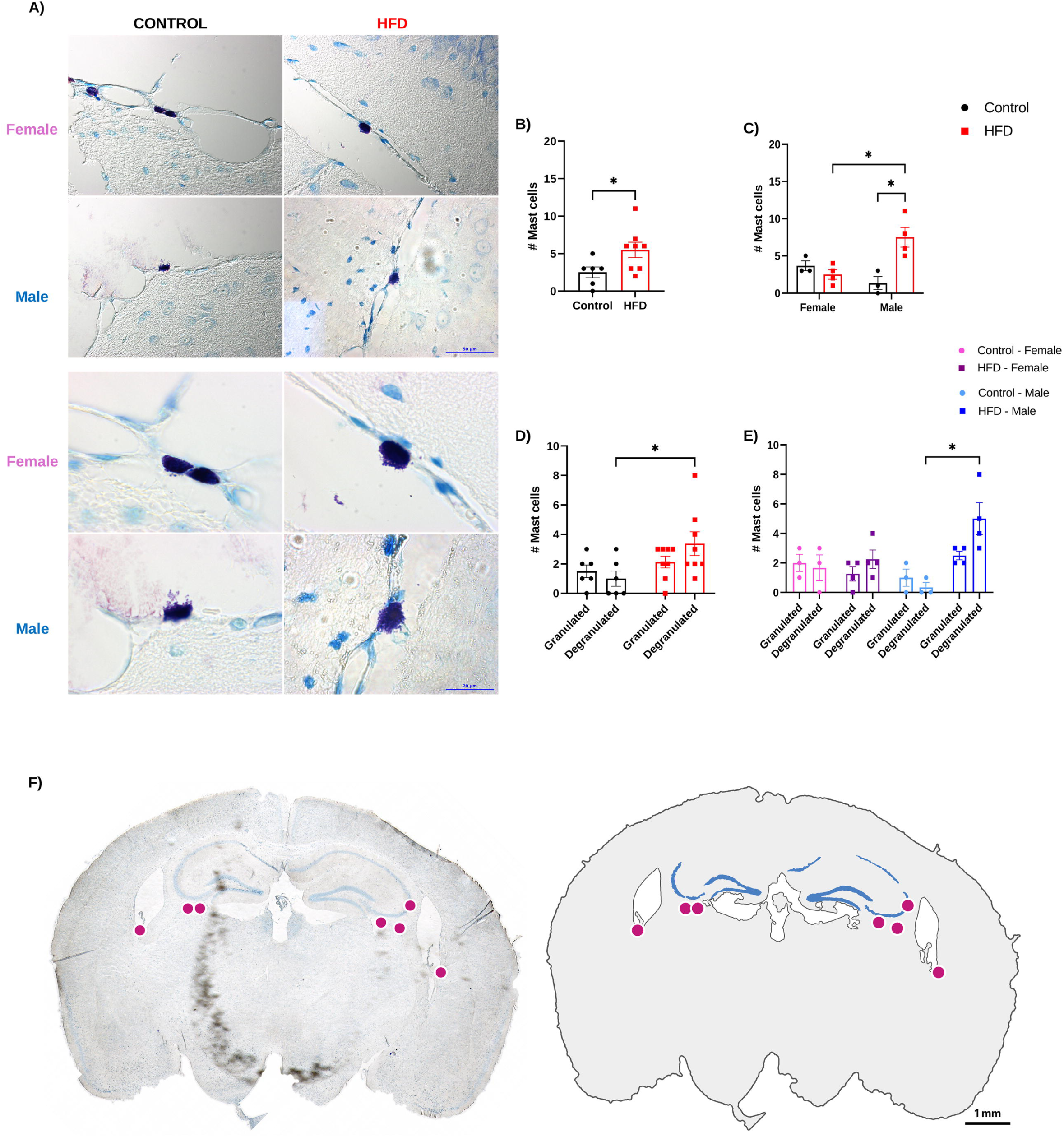
Mast cell identification by toluidine blue staining in the hippocampal region of mice fed a control or high-fat diet. (A) Representative images of toluidine blue-stained coronal brain sections showing the hippocampus in different groups (40x; scale bar = 20 µm). The graphs show differences in the number of mast cells in the hippocampus and adjacent regions, including the lateral ventricles, the third ventricle, the corpus callosum, and the part of the cortex near the hippocampus (hippocampal region), in mice fed a control diet or a high-fat diet for 16 weeks (B), as well as differences by sex (C). Analysis of the number of granulated or degranulated mast cells in the hippocampal region of mice fed a control diet and a high-fat diet (D), and differences by sex (E). The number of mast cells was determined by analyzing two slides from each mouse. F) Localization of mast cells in the hippocampal region. Left, a coronal brain section at the level of the dorsal hippocampus from a high-fat diet-fed male mouse, stained with toluidine blue and reconstructed from 38 overlapping 5× fields. Right, a schematic tracing of the same section showing the lateral ventricles and the hippocampal cell layers (blue). Magenta dots mark the positions of mast cells identified in this section. Mast cells were found in the lateral ventricles and along the ventral border of the hippocampus. Scale bar, 1 mm. The data were analyzed using an unpaired t-test to assess differences between the control and high-fat diet groups. Sex-dependent differences, as well as the numbers of granulated and degranulated mast cells, were analyzed using a two- or three-way ANOVA, followed by a Tukey post hoc test. The data are presented as mean ± SEM (* p < 0.05).

When comparing the numbers of granulated and degranulated mast cells, the high-fat diet group had more degranulated mast cells than the control diet group (Fig. 6D; *p* = 0.0436). Notably, only male mice fed a high-fat diet showed an increase in degranulated mast cells compared with those fed a control diet (Fig. 6E; *p* = 0.0021; interaction diet/sex *p* = 0.0037).

### 3.6 A high-fat diet increases mast cell counts and triggers degranulation in the superficial cervical lymph nodes of male mice

Because mast cells reside in the superficial cervical lymph nodes, we counted both total and degranulated mast cells in these lymph nodes. We also examined the inguinal lymph nodes, which are anatomically distant from the brain.

Overall, the control and high-fat diet groups showed similar total numbers of mast cells in the superficial cervical lymph nodes (Fig. 7C). However, when analyzing the data by sex, female mice fed a high-fat diet had fewer mast cells (Fig. 7D; *p* = 0.0324), with a significant interaction between diet and sex (*p* = 0.0019). In contrast, male mice fed a high-fat diet tended to have more mast cells than their control group, but the difference was not significant (*p* = 0.1102). Relative to female mice fed a high-fat diet, male mice on this diet showed a significant increase in the number of mast cells (Fig. 7D; *p* = 0.0456). To evaluate the spatial distribution of mast cells within superficial cervical lymph nodes, we performed automated whole-node mapping and quadrat-based spatial pattern analysis (Fig. 7A). Reconstructed whole-node sections revealed that mast cells were not uniformly distributed; rather, the vast majority localized preferentially to the perinodal and capsular periphery, with significantly fewer cells scattered throughout the dense lymphoid parenchyma.

**Figure 7.**
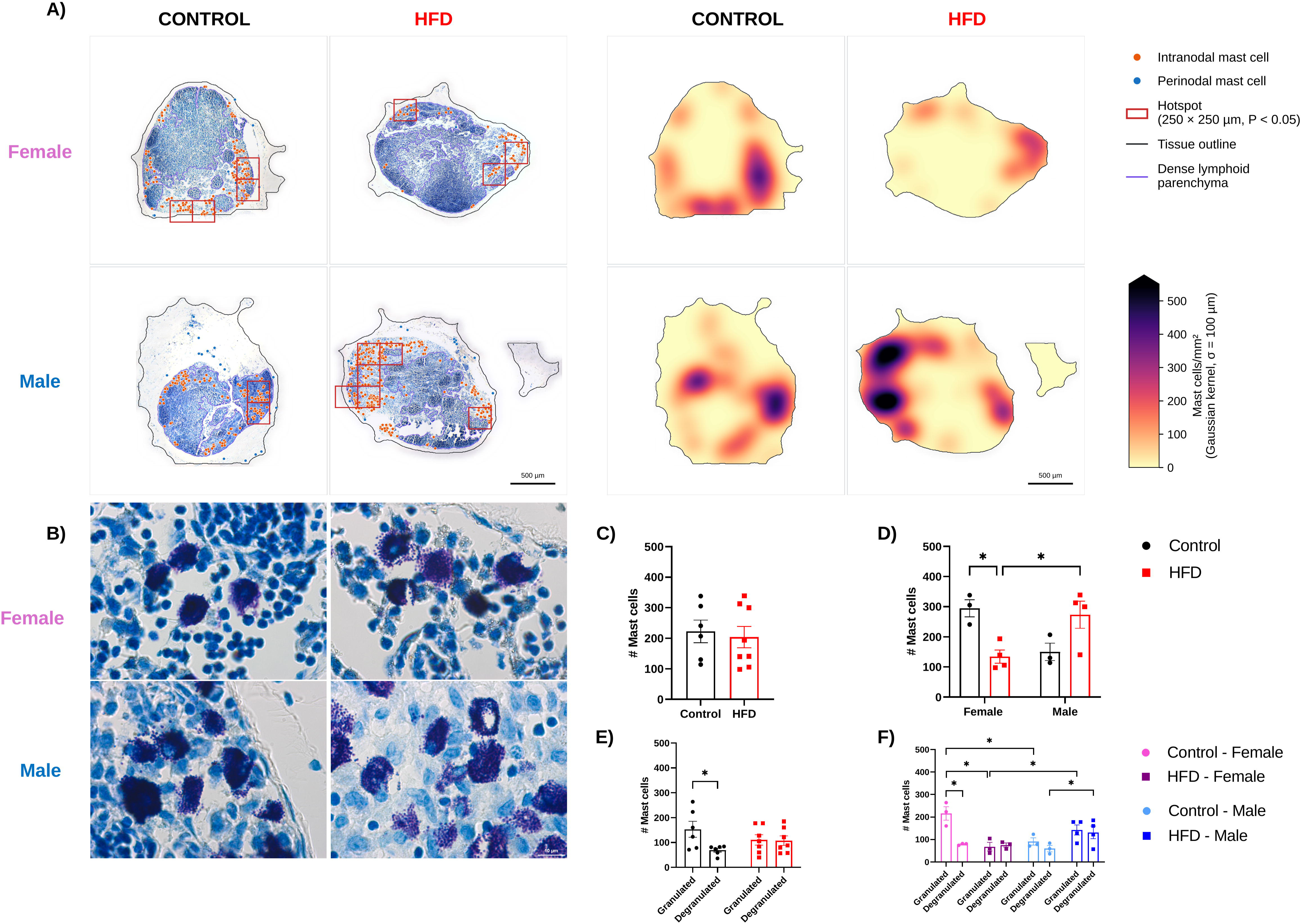
Mast cells in superficial cervical lymph nodes of mice fed a control or high-fat diet identified by toluidine blue staining. (A) Representative superficial cervical lymph nodes from control and high-fat diet fed females and males mice. Left, whole lymph node sections reconstructed from 5× fields, showing the position of each mast cell (orange, intranodal; blue, perinodal), the tissue outline (black), the dense lymphoid parenchyma (purple), and mast cell hotspots (red squares; 250 × 250 µm quadrats with more mast cells than expected under complete spatial randomness, P < 0.05). Right, local mast cell density in the same sections (mast cells/mm², Gaussian kernel, σ = 100 µm), shown on the same color scale for all panels. Mast cells were detected automatically in 20× images and mapped onto the reconstructions. Scale bars, 500 µm. (B) Toluidine blue staining at 100× in control and high-fat diet females and males; mast cells appear purple. Scale bar, 10 µm. (C) Number of mast cells in superficial cervical lymph nodes of mice fed a control diet or high-fat diet for 16 weeks, and (D) by sex. (E) Number of granulated and degranulated mast cells in control and high-fat diet mice, and (F) by sex. Mast cells in C–F were counted manually in two lymph nodes from each mouse on a single slide (control, n = 6: 3 females and 3 males; high-fat diet, n = 7: 3 females and 4 males). Differences between diets were analyzed with an unpaired t-test; sex-dependent differences and the numbers of granulated and degranulated mast cells were analyzed with two- or three-way ANOVA followed by Tukey’s post hoc test. Data are presented as mean ± SEM; *p < 0.05.

Next, granulated and degranulated mast cells were evaluated to assess the degree of activation. The control group showed fewer degranulated mast cells (Fig. 7E; *p* = 0.0140), whereas the high-fat diet group had equal numbers of granulated and degranulated mast cells (*p* = 0.9188). When analyzing the data by sex, female mice fed a high-fat diet had fewer granulated mast cells than those in the control group (Fig. 7F; *p* = 0.0035) and than male mice fed a high-fat diet (*p* = 0.0217). The data also revealed a significant diet-sex interaction (*p* = 0.0003). Furthermore, high-fat diet consumption in male mice increased degranulated mast cells compared with their control group (Fig. 7F; *p* = 0.0252).

Finally, the inguinal lymph nodes showed no significant changes in mast cell numbers or degranulation (Supplementary Fig. 3A-D), indicating that the high-fat diet increases mast cell numbers and degranulation only in the superficial cervical lymph nodes of male mice.

### 3.7 High-fat diet promotes an inflammatory state in the superficial cervical lymph nodes

Given the increase in the number of mast cells in the superficial cervical lymph nodes of male mice fed a high-fat diet, we then evaluated the relative mRNA expression levels of multiple inflammatory mediators to determine whether the superficial cervical lymph nodes exhibit a specific inflammatory profile (Supplementary Fig. 4). The high-fat diet group showed increased relative expression of the *P2X4* purinergic receptor (*p* = 0.04); the cytokines *IL-1*β (*p* = 0.0192) and *TNF-*α (*p* = 0.04); the integrins *ITGAL* (*p* = 0.0041) and *ITGA4* (*p* = 0.0473); and the chemokine receptors *CCR1* (*p* = 0.0281), *CCR2* (*p* = 0.0052), and *CCR5* (*p* = 0.0054) compared with controls. When we analyzed the data by sex, the superficial cervical lymph nodes of male mice fed a high-fat diet showed increased relative expression of the cytokine *IL-1*β (*p* = 0.01), the integrin *ITGAL* (*p* = 0.0499), and the chemokine receptor *CCR1* (*p* = 0.0106) compared with controls. Compared with female mice fed a high-fat diet, male mice showed increased relative expression of the *P2X1* (*p* = 0.0285), *P2X4* (*p* = 0.027), and *P2X7* (*p* = 0.0259) purinergic receptors; the *NLRP3* inflammasome component (*p* = 0.0497); the cytokine *IL-1*β (*p* = 0.0123); and the chemokine receptors *CCR1* (*p* = 0.0015), *CCR5* (*p* = 0.0189), and *CCR7* (*p* = 0.046) in the superficial cervical lymph nodes. In all analyses, the *p* -value for the diet-sex interaction was > 0.06.

Because we observed increased relative CCL3 expression in the hippocampus but no change in CCL3 protein levels, we tested whether lymphatic responsiveness to this chemokine was altered by quantifying the relative expression of its receptors, CCR1 and CCR5, in superficial cervical lymph nodes. Although both receptors showed an increase in their relative expression, CCL3 protein content showed a trend toward increase in males that consumed the high-fat diet (Control-female: 1313 ± 1005; Control-male: 2533 ± 353.1; high-fat diet-female: 252.7 ± 79.90; high-fat diet-male: 5563 ± 2870; p > 0.9999), but it was not statistically significant (Supplementary Fig. 5).

We conducted a correlation analysis of inflammatory markers in superficial cervical lymph nodes and their associations with blood glucose, cholesterol, and hippocampal function. The matrix showed positive correlations between most inflammatory mediators and metabolic variables, particularly weight gain, blood glucose, and cholesterol. Negative correlations were observed between inflammatory mediators and the discrimination index or exploration of the novel object (Supplementary Fig. 6A).

When analyzing female mice, we observed low correlations between variables and inflammatory mediators, with only purinergic signaling components (P2X1, P2X4, P2X7, NLRP3) and metabolic variables (visceral fat, blood glucose, triglycerides) showing negative correlations (Supplementary Fig. 6B). Male mice data showed stronger positive correlations between inflammatory mediators and metabolic variables, and negative correlations with the discrimination index and novel object exploration (Supplementary Fig. 6C). Only P2X1 and FcER1a expression levels showed little or no correlation.

To determine whether the effects of a high-fat diet changes on superficial cervical lymph nodes, which are closely connected to the brain, were specific rather than systemic, we assessed inflammatory mediator mRNA in inguinal lymph nodes, located far from the brain (Supplementary Fig. 7). Most mediators showed no significant change compared to controls (*p* > 0.1), but *NLRP3* (*p* = 0.0364) and *IL-1*β (*p* = 0.0299) were elevated in high-fat diet mice. *CCR5* was also increased in female mice on a high-fat diet (*p* = 0.0178). In all analyses, the p-value for the diet-sex interaction was > 0.08. These findings suggest that a high-fat diet promotes inflammation in the superficial cervical lymph nodes but not in the inguinal lymph nodes.

### 3.8 Cervical lymphadenectomy prevents harmful effects caused by the consumption of a high-fat diet

To confirm the role of superficial cervical lymph nodes in the development of alterations caused by high-fat diet consumption, we surgically removed them via bilateral cervical lymphadenectomy before initiating the high-fat diet. Lymphadenectomy did not significantly affect food intake. In mice fed the high-fat diet, those that underwent lymphadenectomy ate 2.87 ± 0.47 g/day, whereas high-fat diet mice without surgery ate 3.14 ± 0.26 g/day (*p* = 0.89). Likewise, among mice fed the control diet, animals with surgery ate 4.60 ± 0.32 g/day and those without surgery 4.90 ± 0.52 g/day (*p* > 0.99). The same pattern held when males and females were analyzed separately. Females fed the control diet ate similar amounts with and without lymphadenectomy (4.54 ± 0.46 vs. 4.69 ± 0.58 g/day), as did females fed the high-fat diet (3.01 ± 0.59 vs. 3.24 ± 0.22 g/day; *p* > 0.99 for both comparisons). In males, those on the control diet ate 4.65 ± 0.21 g/day with surgery and 5.07 ± 0.47 g/day without lymphadenectomy, and those on the high-fat diet ate 2.68 ± 0.17 and 3.08 ± 0.28 g/day, respectively (*p* > 0.45 for both comparisons). Surprisingly, mice that underwent superficial cervical lymph node removal showed no weight gain (Fig. 8B; *p* > 0.9999) or visceral fat accumulation (Fig. 8E; *p* > 0.9999) after consuming the high-fat diet. This effect persisted when mice were analyzed by sex, with both female (Fig. 8C and F; *p* > 0.9999) and male (Fig. 8D and G; *p* > 0.9999) mice fed a high-fat diet showing no differences compared with their control groups.

**Figure 8.**
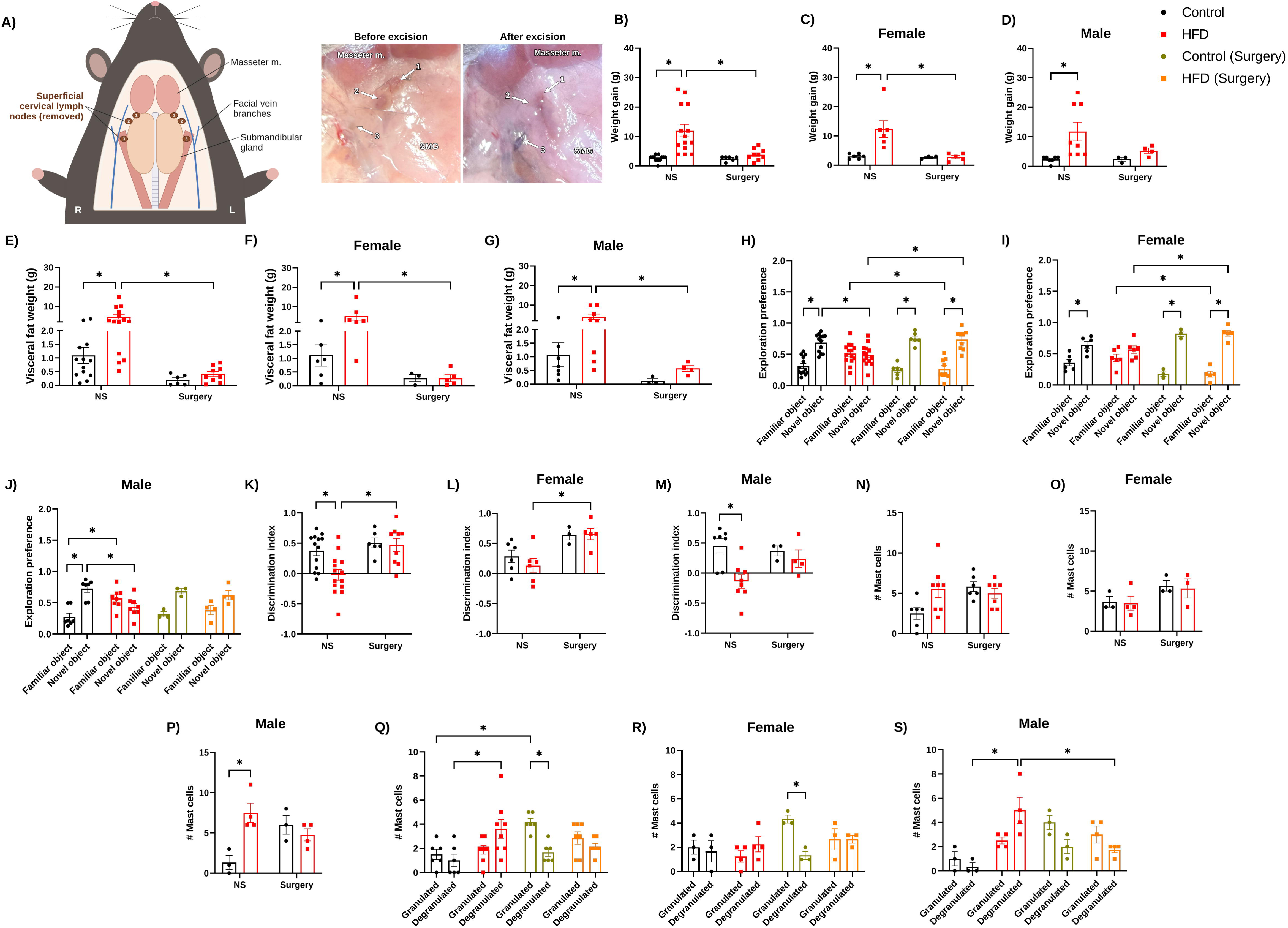
Effect of superficial cervical lymphadenectomy on changes induced by a high-fat diet in female and male mice. (A) Left: Schematic ventral view of a mouse in the supine position with the neck skin elevated, showing the three superficial cervical lymph nodes that have been removed from each side (1–3, brown). These lymph nodes belong to the mandibular group and are located between the masseter muscle and the submandibular gland, near the facial veins. Right: intraoperative photographs of the same surgical field under the stereomicroscope, showing the three lymph nodes before excision and their original locations after excision (arrows, 1–3). Masseter m., masseter muscle; SMG, submandibular gland; R, right; L, left. The graphs show weight gain (B–D), visceral fat accumulation (E–G), exploration preference for a familiar or novel object (H–J), the discrimination index (K–M), the number of mast cells in the hippocampal region (N–P), and the number of granulated and degranulated mast cells (Q–S) in female and male mice without surgery (NS) and with surgery, fed a control diet or a high-fat diet for 16 weeks. Differences between groups were assessed using multiple Mann-Whitney tests and two- or three-way ANOVAs, followed by a Bonferroni post hoc test. The data are presented as mean ± SEM (* p < 0.05).

When mice’s performance in the novel object recognition test was evaluated, those that underwent bilateral superficial cervical lymphadenectomy showed no short-term memory impairment. Regarding exploration time, control mice that did not undergo lymphadenectomy spent more time exploring the novel object than the familiar one (Supplementary Fig. 8A; *p* < 0.0001), whereas mice fed a high-fat diet spent less time exploring the novel object than the control group (*p* < 0.0001). Time spent with the novel object did not differ between high-fat diet-fed mice with and without surgery (*p* > 0.9999). This pattern was similar in females (Supplementary Fig. 8B; *p* > 0.9999) and in males (Supplementary Fig. 8C; *p* > 0.9999). The group fed a high-fat diet showed a significant increase in exploration preference for the novel object compared to their exploration of the familiar objects (Fig. 8H; *p* < 0.0001) and compared with the exploration of the novel object shown by mice fed a high-fat diet but not subjected to surgery (*p* = 0.0066).

Sex-specific analysis revealed that female mice fed a high-fat diet and lacking superficial cervical lymph nodes preferred to explore the novel object compared with the high-fat diet-fed group without surgery (Fig. 8I; *p* = 0.0361). Meanwhile, superficial cervical lymph node resection in male mice fed a high-fat diet prevented the reduction in exploration preference for the novel object observed in male mice fed a high-fat diet without surgery compared with controls (Fig. 8J; *p* = 0.0136). The decreased discrimination index in mice fed a high-fat diet (Fig. 8K; *p* = 0.0083) was restored after bilateral superficial cervical lymphadenectomy (*p* = 0.0026). This effect persisted when the data were analyzed separately for female mice (Fig. 8L; *p* = 0.0168). Meanwhile, male mice fed a high-fat diet without lymphadenectomy showed a reduced index compared with controls (Fig. 8M; *p* = 0.0073). Control mice and those fed a high-fat diet that underwent lymphadenectomy showed a similar discrimination index (p >0.9999), and comparing the high-fat diet groups with and without surgery revealed no changes (p = 0.3206).

Finally, we evaluated the effect of surgery on the number of mast cells near the hippocampus. The no-surgery group fed a high-fat diet showed a trend toward an increase in the number of mast cells, but this was not statistically significant (Fig. 8N; *p* = 0.0840). However, removal of the superficial cervical lymph nodes promoted a similar number of mast cells in the hippocampus-adjacent regions (Fig. 8N). Similar results were observed in female mice, with no differences among groups (Fig. 8O). In male mice, a high-fat diet increased the number of mast cells compared to controls (Fig. 8P; *p* = 0.0104), whereas lymphadenectomy resulted in a similar number of mast cells (*p* > 0.9999).

When classifying mast cells as granulated or degranulated, we found that the number of degranulated mast cells increased in mice fed a high-fat diet compared with controls (Fig. 8Q; *p* = 0.0109), an effect that was prevented by removing the superficial cervical lymph nodes. After stratification by sex, the data revealed that surgery reduced the number of degranulated mast cells in female mice fed the control diet (Fig. 8R; *p* = 0.0496). Meanwhile, male mice fed a high-fat diet showed an increase in the number of degranulated mast cells compared with controls (Fig. 8S; *p* = 0.0014), an effect that was reversed upon removal of the superficial cervical lymph nodes; this was evidenced by a lower number of degranulated mast cells in the group of high-fat diet-fed mice that underwent surgery compared to those that did not undergo surgery (*p* = 0.0276). Overall, these results suggest that superficial cervical lymph nodes play a key role in high-fat diet-induced weight gain and fat accumulation, which in turn prevented the high-fat diet-related changes observed in the novel object recognition test and mast cell activation near the hippocampus.

## 4 Discussion

In this study, we showed that after 16 weeks of feeding a high-fat diet, male mice exhibited impaired short-term memory test performance, whereas female mice were unaffected. In male mice, this vulnerability was associated with inflammatory markers in the superficial cervical lymph nodes, whereas the hippocampus showed only increased CCL3. This finding coincided with increased mast cell numbers and degranulation in the hippocampus and the superficial cervical lymph nodes. The deleterious effects of the high-fat diet on hippocampal function, mast cell number, and degranulation were completely abolished when the superficial cervical lymph nodes were removed, partly because the lymphadenectomy prevented weight gain and fat accumulation. Overall, these results indicate that these cervical lymph nodes and mast cells contribute to hippocampal dysfunction after high-fat diet consumption, with a sex-dependent component in which male mice are more affected than female mice.

The hippocampus is vulnerable to the effects of high-fat diet consumption in male mice, as shown by our results and those of other authors (Hwang et al., 2010; N’Diaye et al., 2024). Reduced neurogenesis markers (Ki67 and DCX) were detected in the hippocampus of male mice fed a high-fat diet (60% of calories from fat) for one month, and these mice also exhibited impaired long-term memory (Baghdadchi et al., 2026). Our results indicate that hippocampal vulnerability is associated with increased relative CCL3 expression. Interestingly, CCL3 might affect hippocampal function by reducing AMPA-dependent neuronal activity, thereby impairing spatial and long-term memory in mice (Marciniak et al., 2015). However, this mechanism may not explain our results, as we did not detect CCL3 protein in the hippocampus. Because weight gain may affect mouse movement, we measured velocity after sixteen weeks of consuming the high-fat diet. Although we did not observe significant changes in speed, future experiments should assess whether the high-fat diet alters motor behavior and the neural circuitry underlying it. On the other hand, evaluating the behavioral performance of mice that consumed the high-fat diet in spatial navigation, working memory, and contextual fear conditioning tests will broaden our understanding of how the high-fat diet affects hippocampal function. Moreover, evaluating electrophysiological activity across different hippocampal neuronal populations after high-fat diet consumption will reveal the underlying electrophysiological changes caused by the high-fat diet. Another potential key contributor to impaired hippocampal activity after sixteen weeks of high-fat diet consumption is astrocytes, as they are active players in glutamatergic synapses (Cuellar-Santoyo et al., 2022).

CCL3 is a chemokine that contributes to the recruitment and infiltration of immune cells such as mast cells, macrophages, monocytes, and neutrophils (Halova et al., 2012; He et al., 2024). We observed an increase in mast cell numbers and degranulation in regions adjacent to the hippocampus of obese male mice. Mast cells are characterized by intracellular granules that contain a variety of modulators, including cytokines, chemokines, proteases, and neurotransmitters. Determining which contents mast cells release in adjacent hippocampal areas would further clarify their contribution to functional alterations in the hippocampus following a high-fat diet. In this sense, using a specific mast cell inhibitor would provide key evidence regarding their contribution to high-fat diet-induced changes. Beyond potentially affecting hippocampal function, mast cells also contribute to obesity development. Mast cell activation of hypothalamic microglia can induce obesity by inhibiting the release of melanocyte-stimulating hormone and altering food intake (Tian et al., 2025). Overall, this information highlights the potential key role of mast cells in brain function and their contribution to impaired hippocampal activity during obesity. Nonetheless, the contributions of other immune cells that respond to CCL3 should be addressed.

The cervical lymph nodes and their resident immune cells may be involved in the brain’s response to chronic systemic inflammation during obesity, given their direct connections to the meninges and choroid plexuses, which allow them to sense and respond to signals originating in the brain and to facilitate the migration of immune cells (Louveau et al., 2018; Rustenhoven et al., 2021; Ruiz-Rodriguez et al., 2026). Our results showed increased mast cells and degranulated mast cells in the superficial cervical lymph nodes of male mice, but not in female mice fed a high-fat diet. Likewise, obese male mice showed increased expression of the inflammatory markers *P2X1*, *P2X4*, *P2X7*, *NLRP3*, *IL-1*β, *ITGAL*, *CCR1*, *CCR5*, and *CCR7* in the superficial cervical lymph nodes, which positively correlated with altered glucose tolerance, increased cholesterol, and impaired short-term memory; a pattern of correlation that was not observed in female mice. Multiple studies in humans and rodents have reported sex-dependent differences in the inflammatory response induced by obesity, with males exhibiting a greater inflammatory response in adipose tissue and lymphoid tissues such as bone marrow, mediated by sex hormones and the expression of genes associated with the sex chromosomes (Singer et al., 2015; Schuetz et al., 2024; Vijayakumar et al., 2026). Our study expands on the mechanism underlying the sex-dependent response during obesity by showing, for the first time, that superficial cervical lymph nodes contribute to obesity-induced inflammation, which underlies the changes in cognitive function we observed only in male mice. This conclusion is also supported by the lack of changes in inflammatory marker expression in the inguinal lymph nodes of obese male mice. These findings suggest that superficial cervical lymph nodes respond to obesity in a sex-specific manner, contributing to the sex dimorphism of obesity-associated inflammation and its effects on hippocampal function.

Elevated inflammatory markers in superficial cervical lymph nodes indicate activation of immune cells, including lymphocytes, neutrophils, monocytes, macrophages, and mast cells, which produce pro-inflammatory cytokines such as IL-1β via purinergic receptor activation (Adinolfi et al., 2026). Increased expression of chemokine receptors and ITGAL might promote migration of immune cells from lymph nodes toward the meninges and, consequently, into the brain (Louveau et al., 2018; Marchetti and Engelhardt, 2020; Shi and Shao, 2023; Lizon et al., 2026). These findings suggest that some immune cells residing in the superficial cervical lymph nodes may migrate toward the meninges or the choroid plexuses, from where they could enter the brain parenchyma. This hypothesis is supported by results from mice with disrupted peripheral-brain communication via deep cervical lymphadenectomy (50). Although the deep cervical lymph nodes have been described as the major drainage route for meningeal lymphatic vessels, these also drain to the superficial cervical nodes (Louveau et al., 2015; Louveau et al., 2018). Jin et al. later demonstrated this contribution. The authors injected a fluorescent tracer into the cisterna magna, and after 120 minutes, the marker was detected in both the superficial and deep cervical lymph nodes (Jin et al., 2025). Interestingly, the percentage of fluorescent tracer accumulation in the deep cervical lymph nodes was about 47%, while the superficial cervical lymph nodes (submandibular (28%) and accessory submandibular (25%) combined accounted for 53% (Jin et al., 2025). To date, deep cervical lymph nodes have been implicated in the pathogenesis of mouse models of multiple sclerosis and glioblastoma (Louveau et al., 2018; Hu et al., 2020; Rustenhoven et al., 2021), but less is known about whether superficial cervical lymph nodes contribute to the pathogenesis of peripheral or brain-related conditions.

Surprisingly, removing superficial cervical lymph nodes prevented high-fat diet-induced weight gain and visceral fat accumulation in both female and male mice. Because male mice on the high-fat diet did not accumulate fat, they did not develop impaired short-term memory and had fewer degranulated mast cells in the hippocampus. These results suggest that superficial cervical lymph nodes play a key role in body fat accumulation in male and female mice, but the exact mechanism remains to be described and may include changes in fat absorption and body metabolism (Kataru et al., 2020). To our knowledge, our research is the first to report that superficial lymph nodes contribute to weight gain and, therefore, to the obesity-related changes we evaluated.

Because we observed increased mast cells in adjacent hippocampal areas and in the superficial cervical lymph nodes in male mice after the high-fat diet, mast cells from the superficial cervical lymph nodes might contribute to the hippocampal alterations seen in male mice. In this sense, genetically altered mast cells or their pharmacological inhibition prevent obesity and glucose intolerance by reducing inflammatory mediators in adipose tissue and acting directly on hypothalamic metabolic regulation (Liu et al., 2009; Tian et al., 2025). However, further studies are needed to confirm whether mast cells in the superficial cervical lymph nodes migrate to the brain or release inflammatory mediators that reach the brain and activate resident mast cells.

## 5 Conclusion

Over sixteen weeks, a high-fat diet impaired short-term memory in male mice, while female mice were not affected. In obese males, the superficial cervical lymph nodes showed increased inflammatory markers, whereas the hippocampus showed only elevated *CCL3* mRNA levels. The hippocampus of obese male mice showed increased total and degranulated mast cells, alongside impaired short-term memory. Therefore, mast cells may contribute to the sex-dependent response to obesity-induced inflammation, which underlies the cognitive dysfunction observed in male mice. Interestingly, removing the superficial cervical lymph nodes prevented body fat accumulation in male and female mice, thereby preventing the deleterious effects of the high-fat diet observed in male mice. Altogether, these results underscore the need to understand the role of superficial cervical lymph nodes in peripheral-brain communication mechanisms, which show a sex-dependent pattern and could guide targeted therapies to prevent obesity and its negative effects on cognitive function.

## Generative AI statement

The author(s) declared that generative AI was not used in the creation of this manuscript. The authors used Claude (Anthropic) to code and debug the image-analysis code. The authors reviewed and validated all outputs and take full responsibility for the content of the publication.

## Author Contributions

**VMR-R:** Conceptualization, Writing – original draft, Writing – review & editing, Formal analysis, Data curation, Methodology, Investigation. **AB-D:** Formal analysis, Methodology, Writing – review & editing. **KH-B:** Formal analysis, Methodology, Writing – review & editing. **AOC-S:** Formal analysis, Methodology, Writing – review & editing. **CMP-P:** Formal analysis, Methodology, Writing – review & editing. **AME-S:** Conceptualization, Project administration, Formal analysis, Writing – original draft, Writing – review & editing, Supervision, Resources, Funding acquisition.

## Acknowledgments

The authors thank the Laboratorio de Microscopía at IPICYT; Dr. Saúl Jijón-Moreno and Dr. J. Sergio Casas-Flores for providing the necessary equipment for the RT-qPCR experiments; and Dr. Roberto Carlos Salgado Delgado for providing materials for tissue staining, as well as Cristian Alonso Galindo-Martínez and Alberto Cardosos Méndez for their help with tissue processing and inclusion.

## Data Availability Statement

The data analyzed in this study are available from the corresponding author upon reasonable request.

