## Supplementary Material for "The superficial cervical lymph nodes and mast cells contribute to sex-dependent cognitive impairment induced by a high-fat diet"

### Supplementary Figures

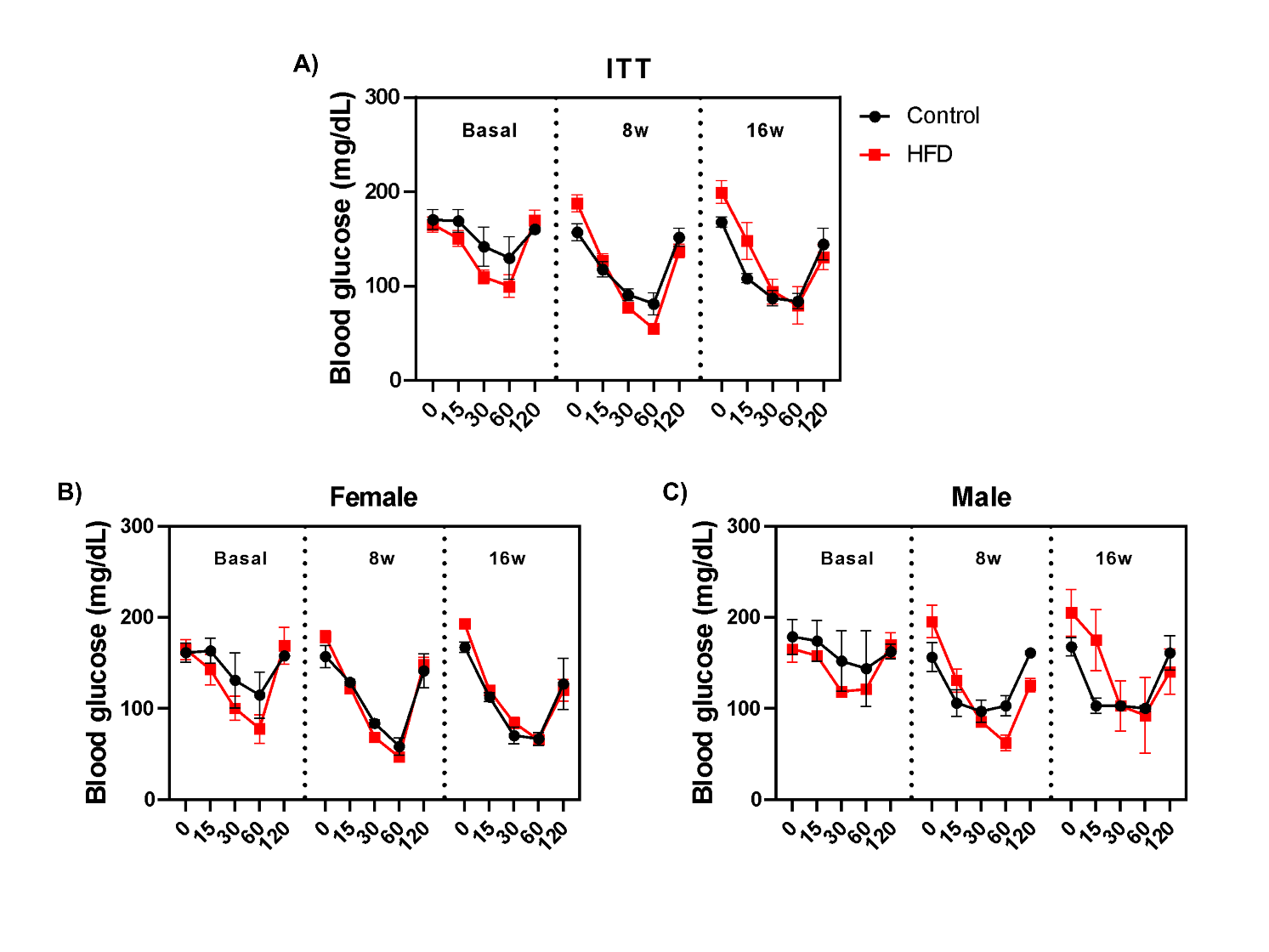

**Supplementary Figure 1. Insulin tolerance test (ITT) in female and male mice fed a control diet or a high-fat diet.** (A) Blood glucose kinetics in mice after an intraperitoneal insulin injection at baseline and at 8 and 16 weeks (8w and 16w) after starting a control diet or a high-fat diet. ITT in female (B) and male (C) mice fed a control diet or a high-fat diet. Data were analyzed using a two-way repeated-measures ANOVA with Bonferroni post hoc testing. Data are presented as mean ± SEM (* p < 0.05).

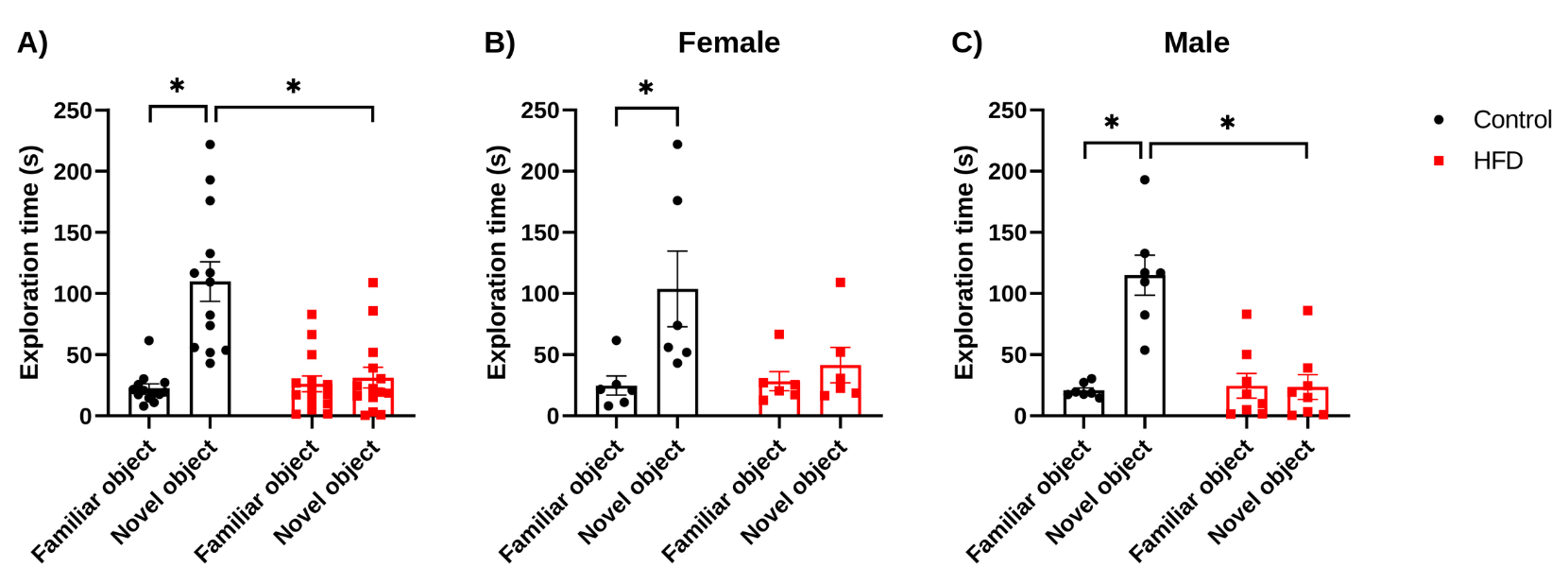

**Supplementary Figure 2. Exploration time in the novel object recognition test in mice fed a control or a high-fat diet.** (A) Time spent exploring the familiar and novel objects by C57BL/6 mice fed a control diet or a high-fat diet during the test session. Exploration time in groups of (B) female and (C) male mice. Data are shown as mean ± SEM, and differences were analyzed using a two-way ANOVA with Bonferroni post hoc testing. *P < 0.05.

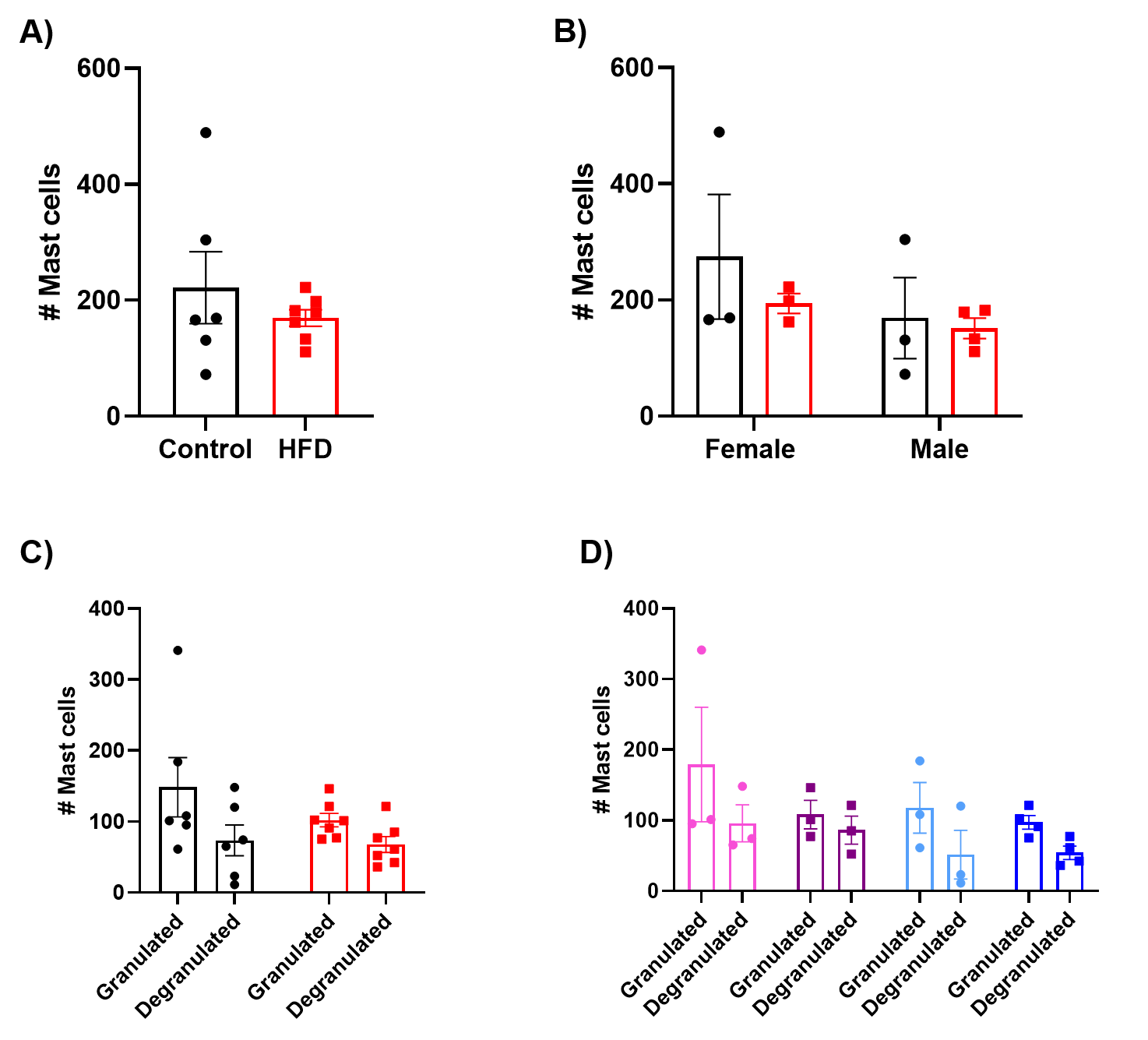

**Supplementary Figure 3. Mast cell identification by toluidine blue staining in inguinal lymph nodes of mice fed a control or high-fat diet.** Assessment of mast cell count (A and B) and degranulation (C and D) in inguinal lymph nodes of female and male mice fed either a control or a high-fat diet. Mast cell count was determined by analyzing two lymph nodes from each mouse on a single slide. Data were analyzed using an unpaired t-test to assess differences between the control and high-fat diet groups. Sex-dependent differences, as well as the numbers of granulated and degranulated mast cells, were analyzed using a two- or three-way ANOVA, followed by a Tukey post hoc test. Data are presented as mean ± SEM (* p < 0.05).

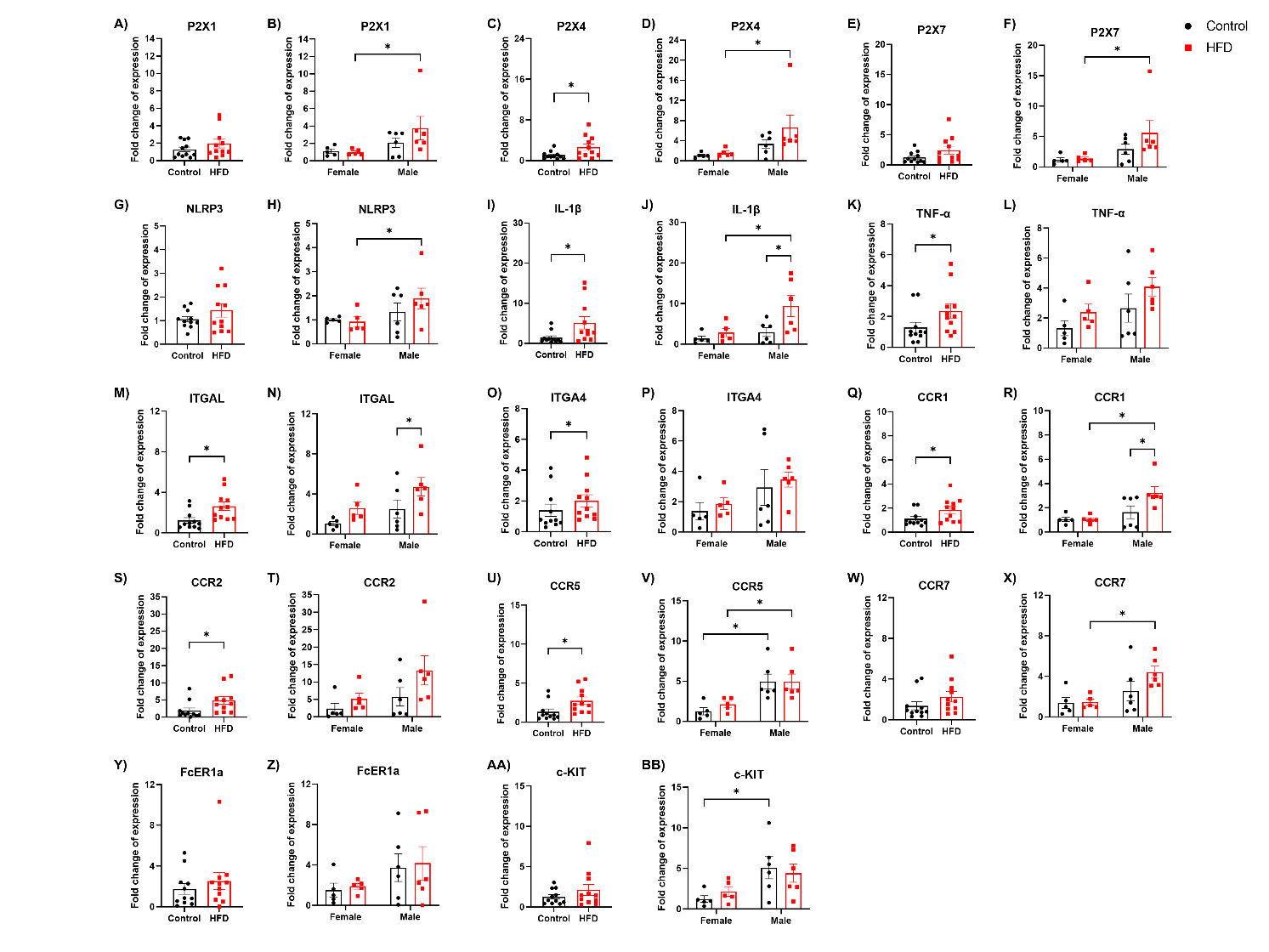

**Supplementary Figure 4. Relative mRNA expression levels of inflammatory mediators in the superficial cervical lymph nodes of mice fed a control or high-fat diet.** The graphs show relative mRNA expression in superficial cervical lymph nodes from mice fed a control or high-fat diet for 16 weeks, with sex differences for the genes encoding the purinergic receptors P2X1 (A and B), P2X4 (C and D), and P2X7 (E and F); the NLRP3 inflammasome component (G and H); the cytokines IL-1β (I and J) and TNF-α (K and L); the integrins ITGAL (M and N) and ITGA4 (O and P); the chemokine receptors CCR1 (Q and R), CCR2 (S and T), CCR5 (U and V), and CCR7 (W and X); and the mast cell markers FcER1a (Y and Z) and c-KIT (AA and BB). Data were analyzed using the unpaired t-test or the Mann-Whitney test to assess differences between the control and high-fat diet groups. To evaluate sex-dependent differences between groups, multiple Mann-Whitney tests or two-way ANOVA with Fisher’s LSD post hoc test were used. Data are presented as mean ± SEM (* p < 0.05).

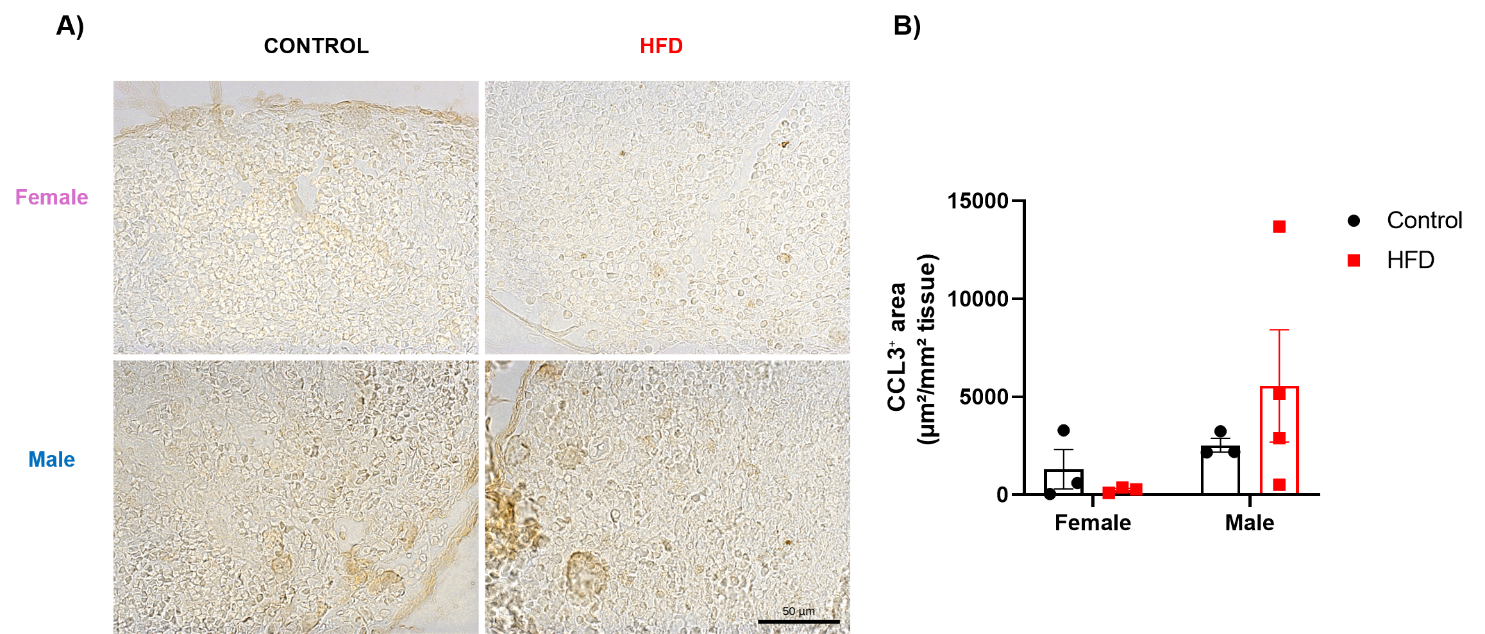

**Supplementary Figure 5. CCL3 immunoreactivity in superficial cervical lymph nodes of control and HFD-fed mice.** (A) Representative bright-field images of CCL3 immunohistochemistry in superficial cervical lymph nodes from female and male mice in the control group and those fed a high-fat diet. Scale bar, 50 µm. (B) Quantification of the CCL3⁺ area, expressed as the DAB-positive area (µm²) per mm² of analyzed tissue. Data are shown as the mean ± SEM (control, n = 3 females and 3 males; HFD, n = 3 females and 4 males).

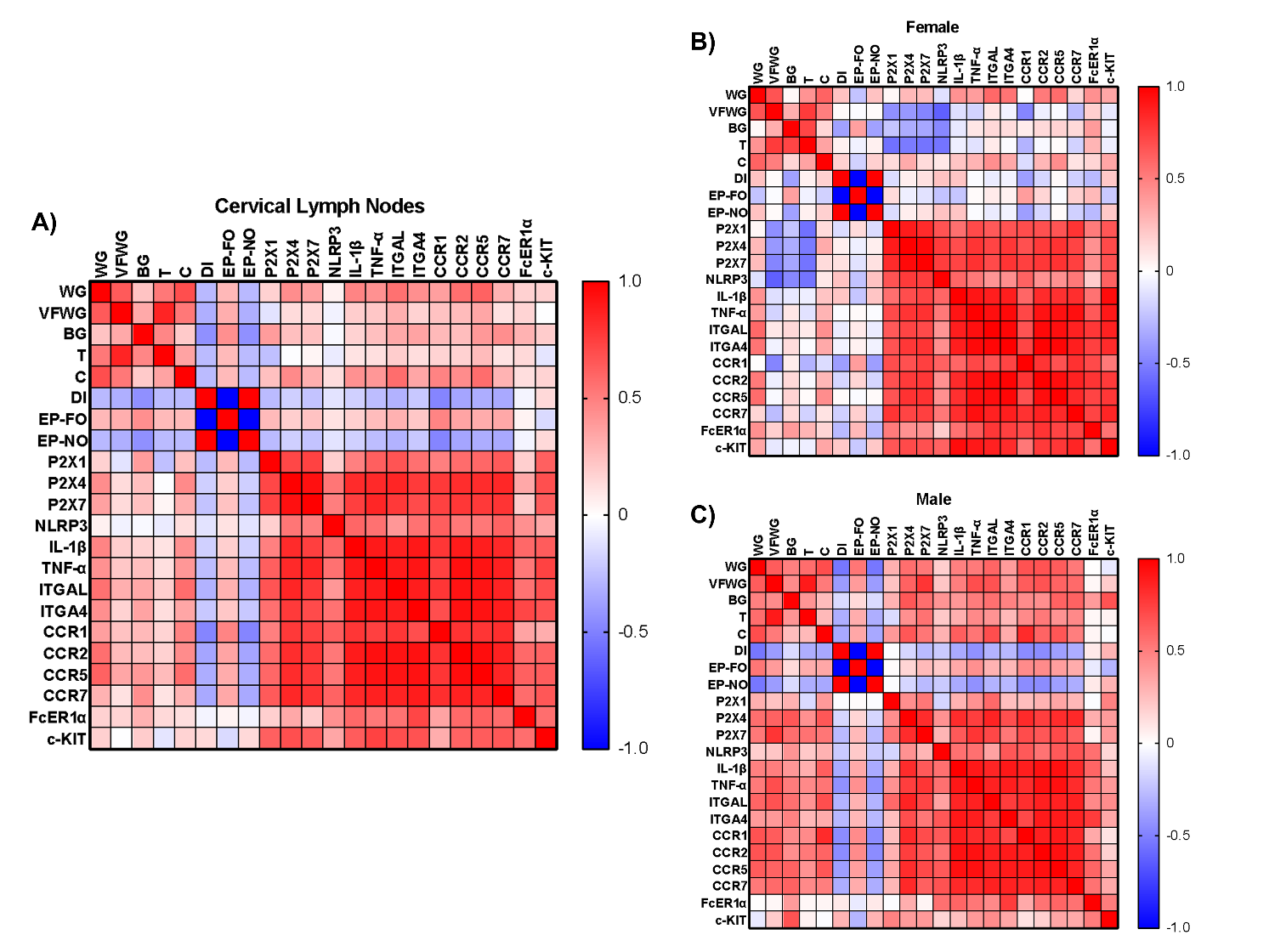

**Supplementary Figure 6. Correlation matrix of variables from cervical lymph nodes of mice fed a control or high-fat diet.** (A) Correlation matrix of inflammatory marker expression levels in cervical lymph nodes with metabolic and cognitive variables in mice fed a control diet or a high-fat diet for 16 weeks. Correlation matrices were obtained by analyzing variables from female (B) and male (C) mice. Data were analyzed using Spearman’s correlation coefficients. Red indicates a positive correlation, blue indicates a negative correlation, and white (0) indicates no correlation. Variables include weight gain (WG), visceral fat weight gain (VFWG), blood glucose (BG), blood triglycerides (T), blood cholesterol (C), discrimination index (DI), exploration preference for the familiar object (EP-FO), and exploration preference for the novel object (EP-NO).

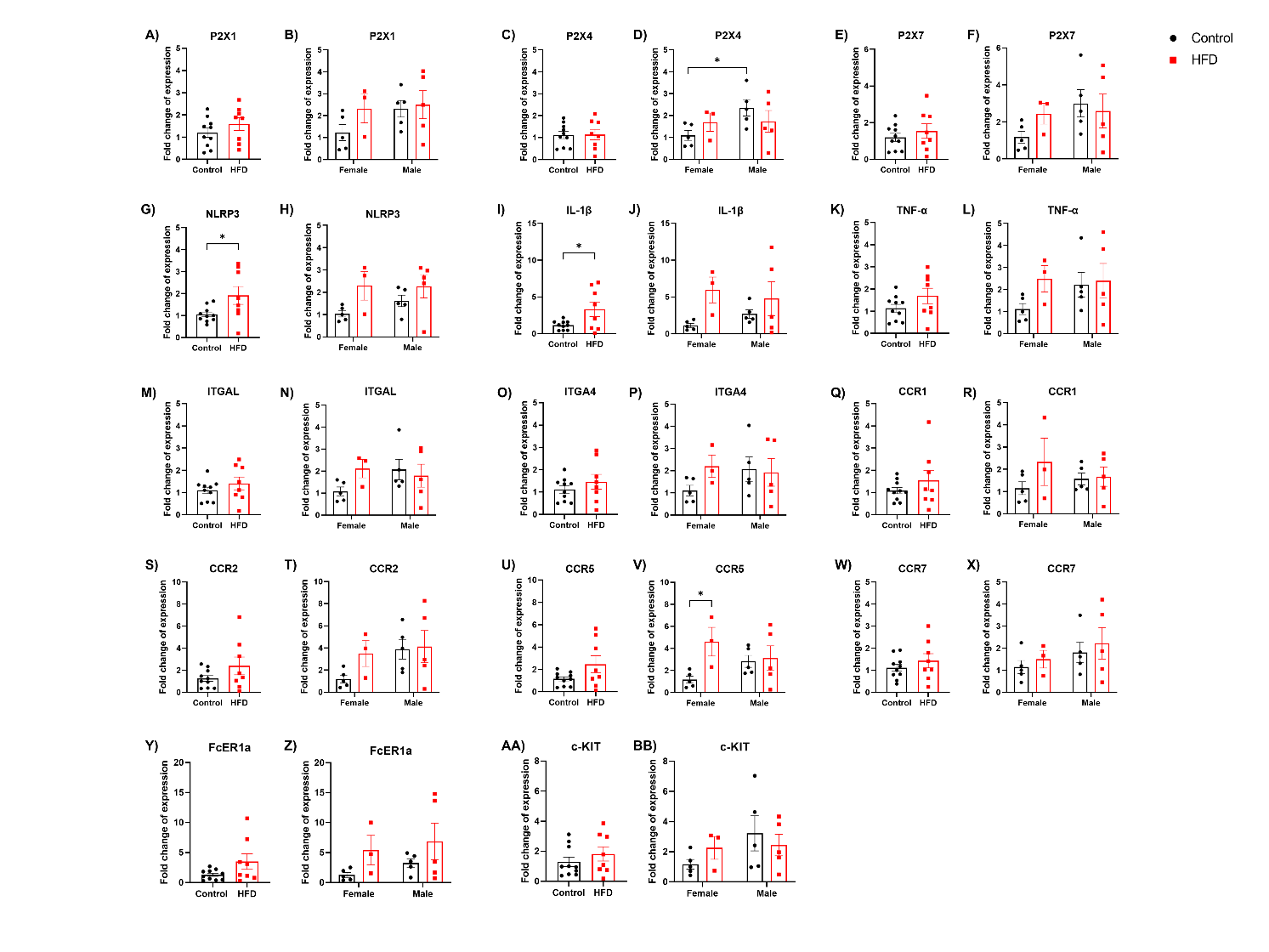

**Supplementary Figure 7. Relative mRNA expression levels of inflammatory mediators in the inguinal lymph nodes of mice fed a control or high-fat diet.** The graphs show relative mRNA expression in inguinal lymph nodes from mice fed a control or high-fat diet for 16 weeks, with sex differences for the genes encoding the purinergic receptors P2X1 (A and B), P2X4 (C and D), and P2X7 (E and F); the NLRP3 inflammasome component (G and H); the cytokines IL-1β (I and J) and TNF-α (K and L); the integrins ITGAL (M and N) and ITGA4 (O and P); the chemokine receptors CCR1 (Q and R), CCR2 (S and T), CCR5 (U and V), and CCR7 (W and X); and the mast cell markers FcER1a (Y and Z) and c-KIT (AA and BB). Data were analyzed using the unpaired t-test or the Mann-Whitney test to assess differences between the control and high-fat diet groups. To evaluate sex-dependent differences between groups, multiple Mann-Whitney tests or two-way ANOVA followed by Fisher’s LSD post hoc test were used. Data are presented as mean ± SEM (* p < 0.05).

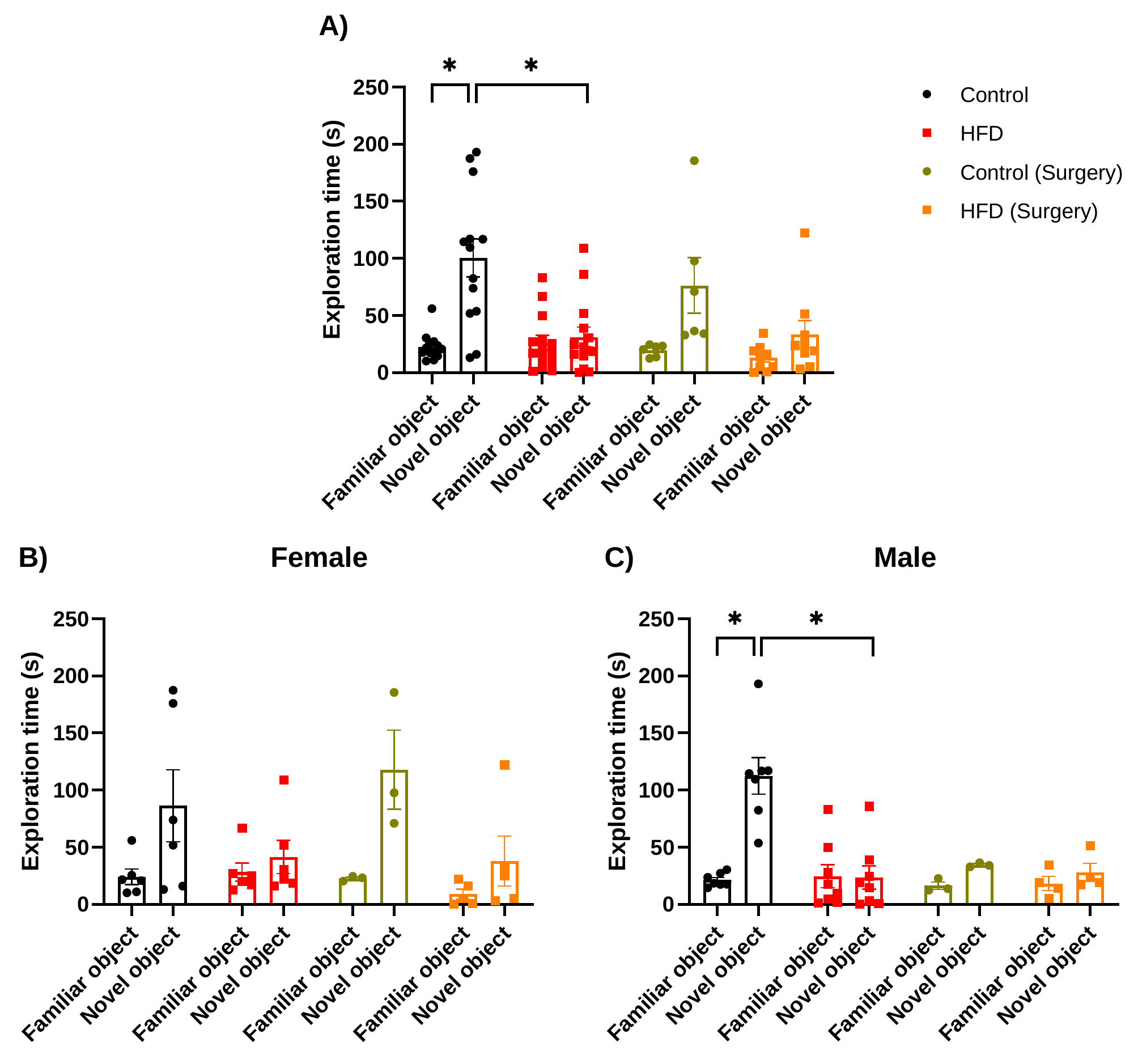

**Supplementary Figure 8. Exploration time in the novel object recognition test in control and mice fed a high-fat diet, with or without lymphadenectomy.** (A) Time spent exploring the familiar and novel objects during the test session in mice fed a control or high-fat diet, with or without surgery. Exploration time for familiar and novel objects in female (B) and male (C) mice. Data are shown as mean ± SEM; differences between groups were analyzed using three-way ANOVAs, followed by Bonferroni post hoc tests (* p < 0.05).
